# Phosphatidylethanolamine facilitates mitochondrial pyruvate entry to regulate metabolic flexibility

**DOI:** 10.1101/2022.11.01.514767

**Authors:** Piyarat Siripoksup, Guoshen Cao, Ahmad A. Cluntun, J. Alan Maschek, Quentinn Pearce, Marisa J. Lang, Hiroaki Eshima, Precious C. Opurum, Ziad S. Mahmassani, Eric B. Taylor, James E. Cox, Micah J. Drummond, Jared Rutter, Katsuhiko Funai

**Affiliations:** Diabetes & Metabolism Research Center, University of Utah, Salt Lake City, Utah, USA; Department of Physical Therapy & Athletic Training, University of Utah, Salt Lake City, Utah, USA; Department of Biochemistry University of Utah, Salt Lake City, Utah, USA; Metabolomics Core Research Facility, University of Utah, Salt Lake City, Utah, USA; Department of Nutrition & Integrative Physiology, University of Utah, Salt Lake City, Utah, USA; Fraternal Order of Eagles Diabetes Research Center, University of Iowa, Iowa City, Iowa, USA; Molecular Medicine Program, University of Utah, Salt Lake City, Utah, USA; Howard Hughes Medical Institute, University of Utah, Salt Lake City, Utah, USA

## Abstract

Carbohydrates and lipids provide the majority of substrates to fuel mitochondrial oxidative phosphorylation (OXPHOS). Metabolic inflexibility, defined as an impaired ability to switch between these fuels, is implicated in a number of metabolic diseases. Here we explore the mechanism by which physical inactivity promotes metabolic inflexibility in skeletal muscle. We developed a mouse model of sedentariness by small mouse cage (SMC) that, unlike other classic models of disuse in mice, faithfully recapitulates metabolic responses that occur in humans. Bioenergetic phenotyping of mitochondria displayed metabolic inflexibility induced by physical inactivity, demonstrated by a reduction in pyruvate-stimulated respiration (*J*O_2_) in absence of a change in palmitate-stimulated *J*O_2_. Pyruvate resistance in these mitochondria was likely driven by a decrease in phosphatidylethanolamine (PE) abundance in the mitochondrial membrane. Reduction in mitochondrial PE by deletion of phosphatidylserine decarboxylase (PSD) was sufficient to induce metabolic inflexibility measured at the whole-body level, as well as at the level of skeletal muscle mitochondria. Low mitochondrial PE was sufficient to increase glucose flux towards lactate. We further implicate that resistance to pyruvate metabolism is due to attenuated mitochondrial entry via mitochondrial pyruvate carrier (MPC). These findings suggest a novel mechanism by which mitochondrial PE directly regulates MPC activity to modulate metabolic flexibility.

## Introduction

Chronic physical inactivity increases all-cause mortality by 30%, accounting for one death every 44 seconds [1–4]. Sedentary behavior exacerbates the risk for many chronic diseases such as type 2 diabetes and cardiovascular diseases [5–7]. Systemic metabolic disturbances induced by inactivity is likely largely responsible for the pathogenesis of these conditions [7, 8]. Described often as “metabolic inflexibility”, long-term sedentariness impairs the ability to switch between glucose and fatty-acids to fuel ATP synthesis [9, 10]. Metabolic inflexibility that occurs with physical inactivity is primarily driven by the suppression of glucose metabolism in skeletal muscle. Disuse likely directly drives the metabolic reprogramming to attenuate glycolytic flux to mitochondria in the absence of elevated energy demand. The mechanism by which skeletal muscle mitochondrial metabolism adapts to chronic disuse is not well understood.

Our understanding of the underlying molecular processes that drive inactivity-induced metabolic inflexibility has been limited partly due to the lack of appropriate pre-clinical models of human sedentary behavior [11]. Traditional murine models of muscle disuse or physical inactivity, such as hindlimb unloading, cast immobilization, and denervation models are well-suited to study muscle atrophy, but they do not phenocopy the systemic and skeletal muscle metabolic adaptations observed in humans [11, 12]. To address this important methodological gap, we adapted a novel mouse model of inactivity, small mouse cage (SMC) [13, 14] that more reliably induces metabolic perturbations with sedentariness. This model now enabled us to more rigorously investigate the interplay between mitochondrial energetics and metabolic inflexibility in the context of physical inactivity.

Previously, we identified mitochondrial phosphatidylethanolamine (PE) to be an important regulator of mitochondrial oxidative phosphorylation (OXPHOS) that is induced by exercise training and suppressed with hindlimb unloading [15]. PE is highly concentrated in the inner mitochondrial membrane (IMM) and is autonomously synthesized by phosphatidylserine carboxylase (PSD) [16, 17]. In mammalian systems, nearly all PE is synthesized in the IMM by PSD and exported to other regions of the cell, while the PE generated by the CDP-ethanolamine pathway in the endoplasmic reticulum does not translocate to mitochondria [18, 19]. Human mutation in the Pisd gene causes mitochondrial disease [20–22]. We have previously shown that skeletal muscle-specific deletion of PSD (homozygous knockout) is lethal due to robust atrophy and weakness of the diaphragm muscle [15]. The consequence of a more modest reduction of mitochondrial PE, such that occurs with sedentariness, is unknown. Importantly, muscle phospholipid composition, particularly low PE, has been linked to metabolic inflexibility in humans [23–26].

In this study, we implicate reduced muscle mitochondrial PE as the driving force behind inactivity-induced metabolic inflexibility. SMC intervention modestly lowered mitochondrial PE, concomitant to the suppression of glucose metabolism. We then recapitulated moderate reductions in mitochondrial PE using a skeletal muscle-specific heterozygous knockout of PSD (PSD-MHet). Unlike their homozygous counterparts, heterozygous deletion of PSD produced modest systemic and skeletal muscle phenotype that resembled many metabolic shifts found with the SMC intervention.

## Results

### SMC housing induces metabolic inflexibility in male but not female mice

Sedentary behavior promotes systemic and skeletal muscle metabolic inflexibility in humans [7, 27]. In contrast, commonly utilized models of disuse in mice such as hindlimb unloading increases skeletal muscle glucose metabolism (Figure 1 – figure supplement 1A). To better model the metabolic disturbances observed in human inactivity, we developed a mouse model of physical inactivity using SMC (Figure 1A). Male and female wild-type C57BL/6J mice were ambulatory or subjected to eight weeks of SMC housing that substantially restricted gross spontaneous movement (Figure 1B). Body mass, lean mass, and individual muscle masses were significantly reduced in male nice and not in female mice (Figure 1C&D and Figure 1 – figure supplement 1B). In contrast, SMC intervention did not alter adiposity in either sex, although there was a trend for greater adipose tissue masses only in female mice (Figure 1E and Figure 1 – figure supplement 1C). To evaluate the effects of reduced activity on metabolic flexibility, mice underwent indirect calorimetry for measurements of whole-body O_2_ consumption (VO_2_) and respiratory exchange ratio [28]. VO_2_ was not influenced with SMC in both sexes (Figure 1F&G), consistent with findings that changes in physical activity do not drive changes in total daily energy expenditure [29]. RER is an indicator of systemic substrate preference, where a value of 1.0 signifies a 100% reliance on carbohydrates, whereas a value of 0.7 indicates a 100% reliance on lipids. Mice rely more on lipids during the light cycle when they are asleep, and shift to carbohydrate utilization during the dark cycle when they are active or eating. Notably, while SMC induced metabolic inflexibility in male mice, female mice demonstrated normal metabolic flexibility (Figure 1H&I). Specifically, SMC reduced the ability of male mice to shift to carbohydrate usage during the dark cycle. Further, consistent with attenuated systemic glucose metabolism, SMC intervention elevated fasting serum glucose in male mice (Figure 1J).

**Figure 1.**
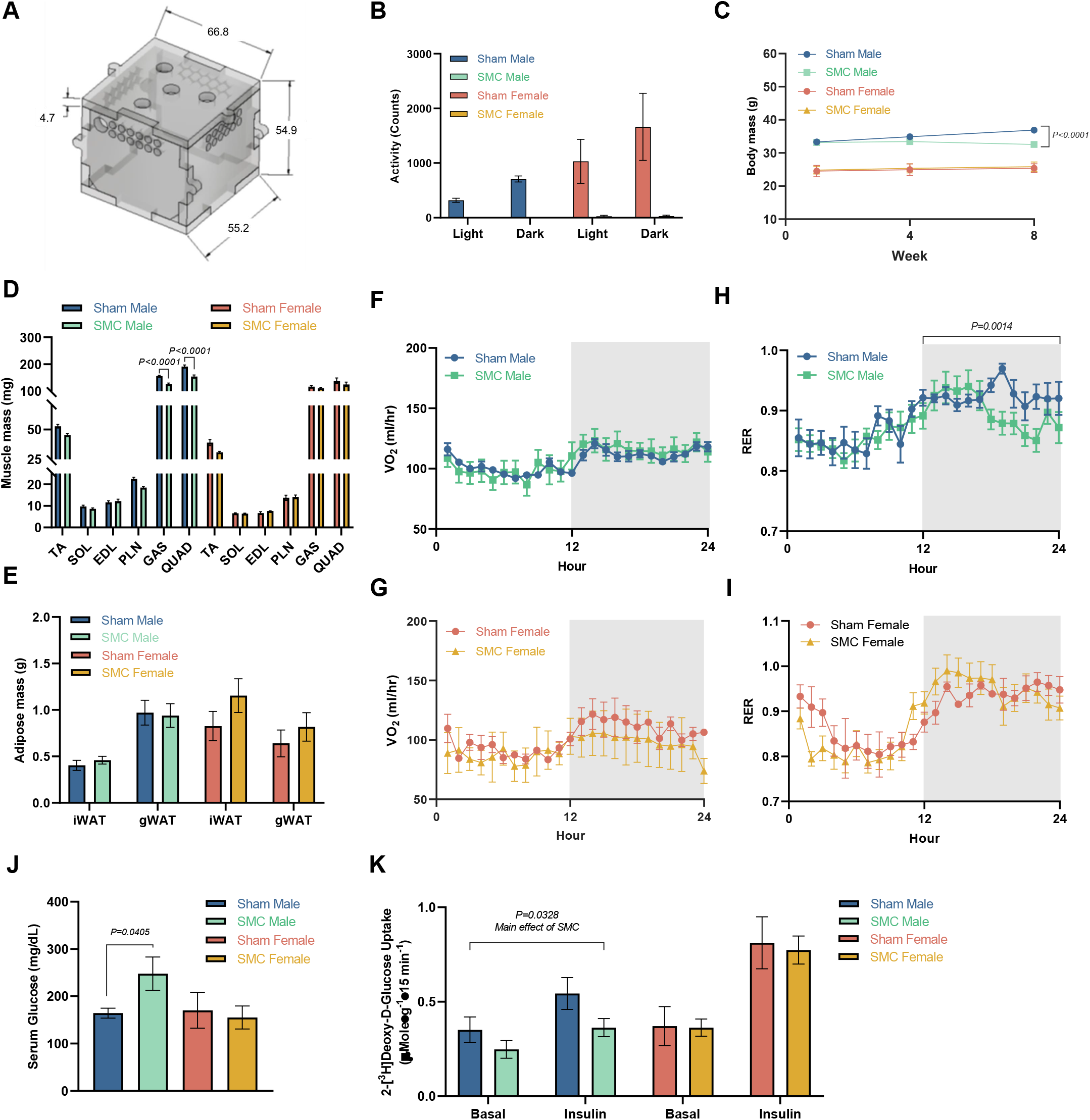
SMC housing induces metabolic inflexibility in male but not female mice. (A) Small mouse cage schematic. (B) Activity counts of sham and SMC mice via indirect calorimetry (n = 8 per group). (C) Body mass time course (n= 6-14 per group). (D) Skeletal muscle tissue mass (N = 7-8 per group). (E) Adipose mass of sham and SMC mice (n = 7-19 per group). (F) Absolute VO_2_ of male sham and SMC mice via indirect calorimetry (n = 8-9 per group). (G) Absolute VO_2_ of female sham and SMC mice via indirect calorimetry (n = 3-4 per group). (H) Respiratory exchange ratio [28] of male sham and SMC mice (n = 8-9 per group). (I) RER of female sham and SMC mice (n = 3-4 per group). (J) Fasting serum glucose levels of sham and SMC mice (n = 4-8 per group). (K) [^3^H]2-deoxyglucose glucose uptake in soleus muscles of male and female sham and SMC mice (n = 4-9 per group). Data represent mean ± SEM. P-values generated by two-tailed, equal variance, Student’s t-test (D and J), or by 2-way ANOVA with Tukey’s post hoc test (B-C, E-I, and J).

To examine glucose metabolism in skeletal muscle, we excised soleus muscles from male and female sham or SMC mice for the measurement of ex vivo 2-deoxyglucose uptake. Congruent with systemic metabolic inflexibility, SMC intervention reduced glucose uptake in both basal and insulin-stimulated conditions in males, but not in females (Figure 1K). These findings are consistent with the hypothesis that reduced skeletal muscle glucose metabolism drives systemic metabolic inflexibility induced by SMC. It is noteworthy that male mice became metabolically inflexible despite no increases in adiposity (Figure 1E and Figure 1 – figure supplement 1C). Metabolic inflexibility also occurred independently of increases in food intake or serum cortisol levels. (Figure 1 – figure supplement 1D&E).

We sought to capitalize on the sexually dimorphic response to explore the mechanism by which SMC induces skeletal muscle metabolic inflexibility only in male mice. RNA sequencing of gastrocnemius muscles followed by KEGG pathway analysis revealed similarities and differences in gene set enrichment in a number of pathways between males and females (Figure 2A). The ribosomal pathway was among the most negatively enriched categories with both sexes, consistent with the notion that inactivity decreases muscle protein synthesis [30]. Notably, metabolic pathways were reduced in males but not in females, suggesting that metabolic reprogramming induced by SMC may be unique to males. Given the central role of mitochondria in these pathways, we further examined the effects of SMC on skeletal muscle mitochondria.

**Figure 2.**
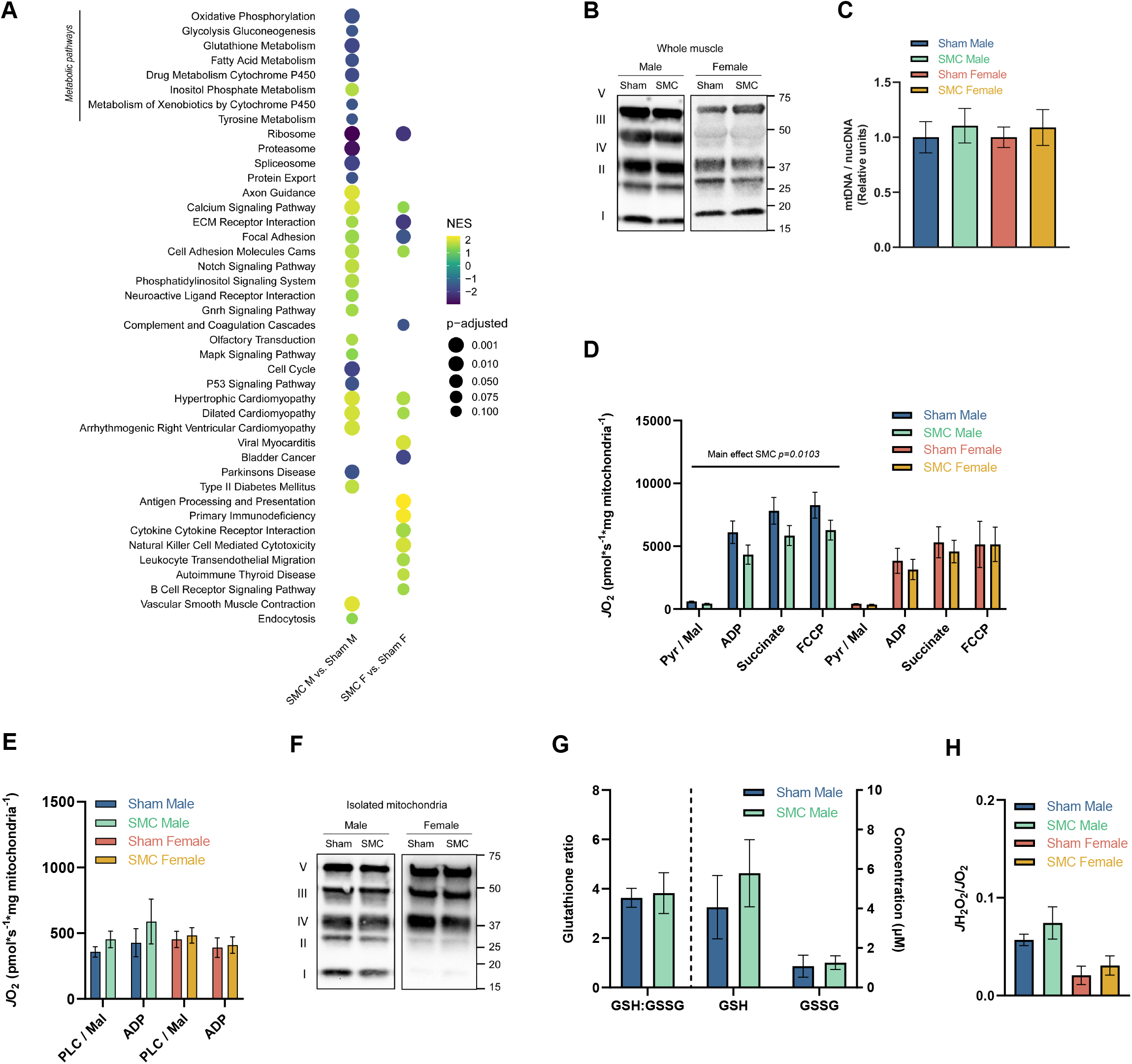
SMC housing reduces pyruvate-dependent respiration without altering palmitate-stimulated respiration. (A) Dot plot representing GSEA pathway analysis (KEGG) of differentially expressing genes (FDR < 0.05) in skeletal muscle of sham and SMC mice. Normalized enrichment scores are represented by a darker color (negatively enriched) and lighter color (positively enriched), while a larger dot diameter indicates a smaller p-adjusted value. Dot plot was generated with R Studio. (B) Representative western blot of ETS protein complexes (I-V) of whole muscle tissue of sham and SMC mice (n = 3-4 per group). (B) Ratio of nuclear to mitochondrial DNA in gastrocnemius muscle (n = 8 per group). (D) O_2_ utilization in isolated mitochondria measured in the presence of 2 mM ADP, 0.5 mM malate, 5 mM pyruvate, 10 mM succinate, 1 μM carbonyl cyanide-*p*-trifluoromethoxyphenylhydrazone (FCCP) of sham and SMC mice (n = 4-6 per group). (E) O_2_ utilization in isolated mitochondria measured in the presence of 2 mM ADP, 0.5 mM malate, 0.02 mM palmitoyl-carnitine (n = 4-6 per group). (F) Representative western blot of ETS proteins in isolated muscle mitochondria of sham and SMC mice (n = 5-6 per group). (G) Glutathione levels in skeletal muscle of sham and SMC mice (n = 6 per group). (H) Electron leak (*J*H_2_O_2_ / O_2_) with succinate in isolated muscle mitochondria from sham and SMC mice (n = 3-4 per group). Data represent mean ± SEM. P-values generated by 2-way ANOVA with Tukey’s post hoc test (C-E, G, and H).

### SMC housing reduces pyruvate-dependent respiration without altering palmitate-stimulated respiration

Previous reports suggest that reduced muscle mitochondrial content can potentially drive metabolic inflexibility induced by inactivity [7, 31]. However, our SMC intervention did not alter mitochondrial density in skeletal muscle (Figure 2B&C and Figure 2 – figure supplement 1A), indicating that lower mitochondrial content is not necessary for inactivity-induced suppression of skeletal muscle glucose metabolism [32]. To this end, we further examined respiratory function per unit of mitochondria. High-resolution respirometry experiments showed that SMC robustly diminished respiration (*J*O_2_) driven by pyruvate in male, but not female mice (Figure 2D), consistent with the notion that metabolic inflexibility is driven by mitochondria’s ability to accept glycolytic substrates. Strikingly, there was no difference in *J*O_2_ fueled by palmitate (Figure 2E), indicating that the reduced ability of mitochondria to accept substrates is limited to glycolytic substrates. Moreover, these changes occurred independently of changes in OXPHOS protein abundance per unit of mitochondria (Figure 2F and Figure 2 – figure supplement 1B).

Some studies indicate that mitochondrial electron leak can promote oxidative stress to suppress glucose metabolism [33]. Multiple labs including our group have reported that traditional models of disuse promote oxidative stress in skeletal muscle [34, 35]. However, our SMC intervention did not alter the ratio of reduced to oxidized glutathione (GSH:GSSG) (Figure 2G) nor reactive lipid aldehydes such as 4-hydroxynonenal (4-HNE) (Figure 2 – figure supplement 1C&D), demonstrating that physical inactivity induced by SMC does not promote oxidative stress. Using high-resolution fluorometry, we further confirmed mitochondrial electron leak (*J*H_2_O_2_/*J*O_2_) to be unaltered with the SMC intervention (Figure 2H). These findings are consistent with results from human bed rest studies [7, 36], ruling out oxidative stress as a mechanism by which SMC intervention suppresses skeletal muscle glucose metabolism.

What is the mechanism by which physical inactivity selectively suppresses mitochondrial pyruvate metabolism in skeletal muscle? SMC intervention had no effect on mRNA levels of pyruvate/glucose metabolism and TCA cycle, nor on protein levels of enzymes of pyruvate metabolism (Figure 3A-C and Figure 3 – figure supplement 1A), indicating that reductions in pyruvate oxidation cannot be attributed to changes in these enzymes. Mitochondrial membrane lipids are known to alter the activity of mitochondrial enzymes in multiple tissues including skeletal muscle [15, 37]. Particularly, disuse induced by hindlimb unloading reduces mitochondrial PE in skeletal muscle [15]. Thus, we examined the effect of SMC housing on the skeletal muscle mitochondrial lipidome. Using LC-MS/MS, we quantified a total of 243 lipids from skeletal muscle mitochondria of sham and SMC mice. Analyses of these lipids revealed a trend for an overall reduction in total phospholipid abundance with SMC in males but not in females (Figure 3 – figure supplement 1B). 73 out of the 243 lipids were significantly downregulated with SMC in male mice (zero upregulated lipids) (Figure 3 – figure supplement 1C) while only two reached statistical significance in female mice (Figure 3 – figure supplement 1D). Among these lipids, PE was most robustly disproportionately downregulated in male mice (Figure 3C&D), consistent with our previous findings with that of hindlimb unloading [15]. Reduced PE with SMC was specific to mitochondria and not reflected in total cellular PE content (Figure 3 – figure supplement 1E&F). Mitochondrial PE is almost exclusively generated by the enzyme PSD, which was substantially reduced in skeletal muscle with SMC (Figure 3E). Thus, we proceeded to investigate the role that mitochondrial PE may play in metabolic inflexibility induced by physical inactivity.

**Figure 3.**
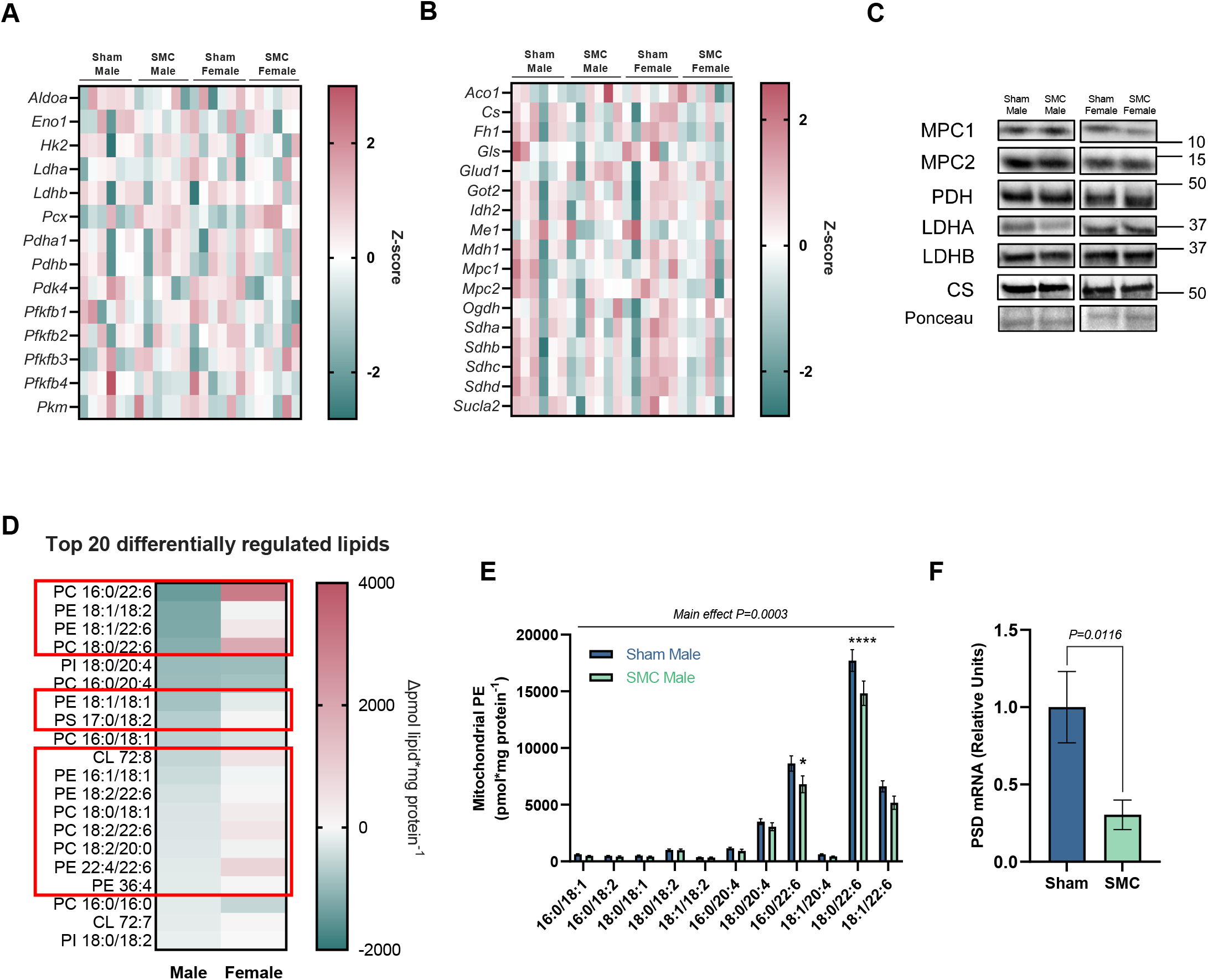
Physical inactivity by SMC housing alters skeletal muscle membrane lipid composition. (A) Heat map of glycolytic genes in sham and SMC mice (n = 6 per group). (B) Heat map of TCA cycle genes in sham and SMC mice (n = 6 per group). (C) Representative western blots of glycolytic/TCA genes in sham and SMC mice (n = 2-6 per group). (D) Top 20 differentially regulated skeletal muscle mitochondrial lipids between SMC and sham mice (n = 7-8 per group). The red box highlights the lipids whose change in abundances are unique to male mice. (E) Skeletal muscle mitochondrial PE species of sham and SMC mice (n = 8 per group). (F) Skeletal muscle PSD mRNA levels of sham and SMC mice (n = 7-8 per group). Data represent mean ± SEM. P-values generated by two-tailed, equal variance, Student’s t-test (F), or by 2-way ANOVA with Tukey’s post hoc test (A-B and D-E).

### Muscle PSD haploinsufficiency makes mice more susceptible to inactivity-induced metabolic inflexibility

Previously, we demonstrated that homozygous deletion of muscle PSD causes lethality due to metabolic and contractile failure in the diaphragm muscle [15]. Homozygous deletion promotes a reduction in mitochondrial PE that is far more robust in magnitude compared to changes in mitochondrial PE observed with SMC. To model a more modest reduction in skeletal muscle mitochondrial PE, we studied mice with tamoxifen-inducible muscle-specific PSD heterozygous deletion (PSD-MHet; PSD^fl/fl^ and HSA-MerCreMer^+/-^) (Figure 4A). As designed, skeletal muscle from PSD-MHet mice had reduced PSD mRNA abundance compared to controls (PSD-MHet; PSD^fl/fl^ and HSA-MerCreMer^-/-^) (Figure 4B), as well as modestly diminished skeletal muscle mitochondrial PE levels (Figure 4C). Unlike the PSD homozygous knockout mice, PSD-MHet appeared normal and healthy under unstressed conditions.

**Figure 4.**
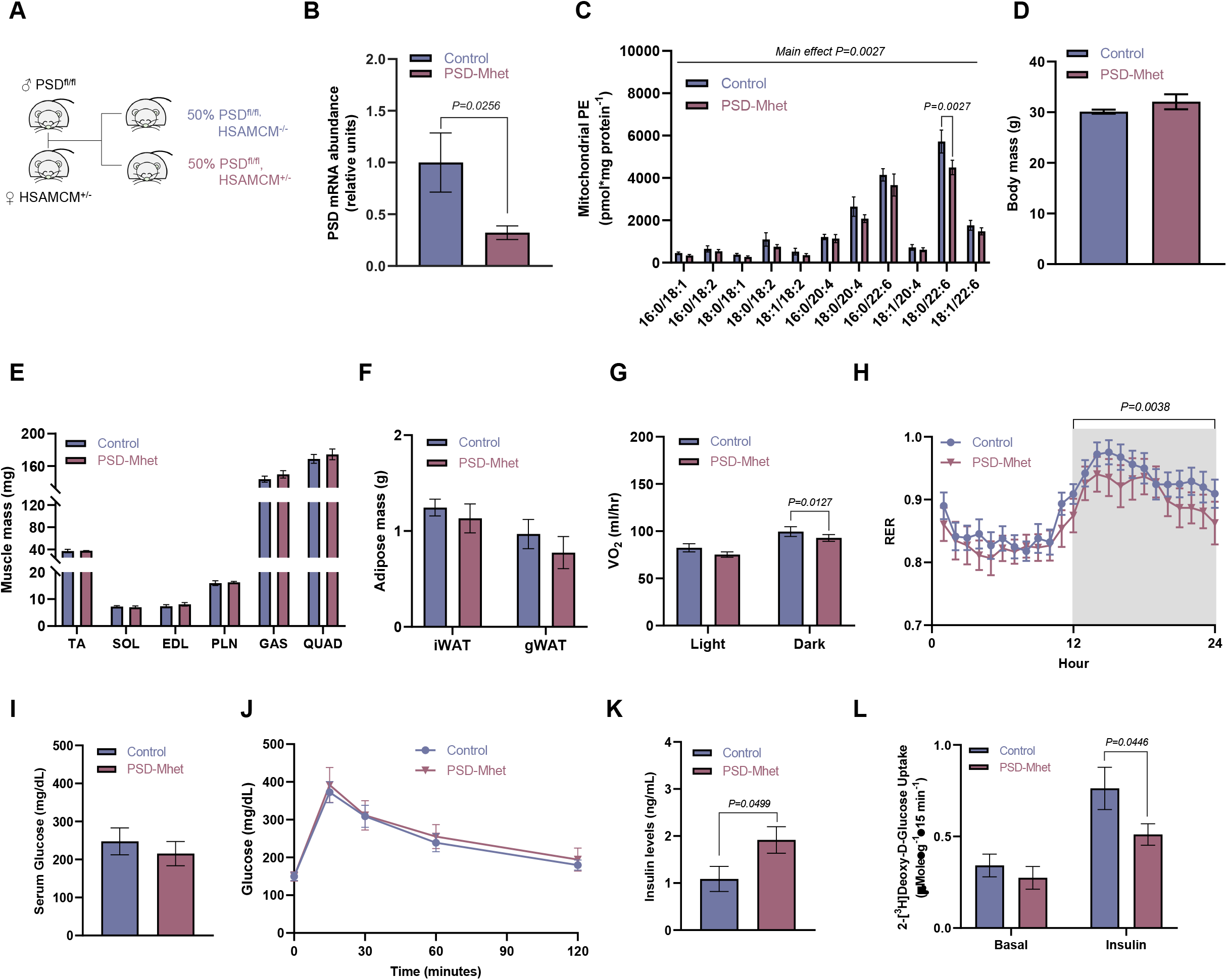
Muscle PSD haploinsufficiency increases susceptibility of mice to inactivity-induced metabolic inflexibility. (A) Mouse breeding schematic. (B) PSD mRNA levels of sham Cre control and PSD-Mhet mice (n = 11-12 per group). (C) Muscle mitochondrial PE levels in sham control and PSD-Mhet mice (n = 5 per group). (D) Body mass of SMC Control and SMC PSD-Mhet mice after 8 weeks of reduced activity (n = 8-12 per group). Skeletal muscle (E) and adipose masses (F) after SMC intervention (n = 8-12 per group). (G) Absolute VO_2_ via indirect calorimetry (n = 3-6 per group). (H) RER (n = 8-11 per group). (I) Serum glucose levels (n = 8 per group). (J) Glucose tolerance test (GTT) performed around Week 7 of SMC intervention (n = 8-13 per group). (K) Serum insulin levels taken at the 30-minute time point during the GTT (n = 6 per group). (l) [^3^H]2-deoxyglucose glucose uptake in soleus muscles after 8 weeks of SMC (n = 7-9 per group). Data represent mean ± SEM. *P < 0.05. **P < 0.01, ***P < 0.001, and ****P < 0.0001. P-values generated by two-tailed, equal variance, Student’s t-test (B, D, I, and K), or by 2-way ANOVA with Tukey’s post hoc test (C, E-H, J, and L).

We placed control and PSD-MHet male mice on eight weeks of SMC to study their systemic and skeletal muscle metabolism. Muscle PSD haploinsufficiency did not influence body mass, body composition, food intake, serum cortisol, or masses of skeletal muscle and adipose tissues (Figure 4D-F and Figure 4 – figure supplement 1A-C). Indirect calorimetry of these mice showed a slight reduction in whole-body VO_2_ in PSD-MHet compared to controls (Figure 4G), which was not explained by changes in physical activity (both virtually undetectably low with SMC). Consistent with our hypothesis that low mitochondrial PE may drive metabolic inflexibility, RER data revealed suppression of glucose metabolism during dark cycle in PSD-MHet mice compared to control mice (Figure 4H). Neither fasting glucose nor glucose tolerance was different between the groups (Figure 4I&J). However, circulating insulin levels at the 30-minute timepoint of the glucose tolerance test was higher in PSD-MHet compared to controls (Figure 4K), suggesting that PSD haploinsufficiency may require greater circulating insulin to stimulate muscle glucose metabolism. Indeed, skeletal muscle glucose uptake was attenuated in PSD-MHet mice compared to control mice (Figure 4L). Collectively, these results suggest that muscle PE deficiency may impair skeletal muscle glucose metabolism to promote metabolic inflexibility.

Similar to our results with the SMC intervention in wildtype mice, PSD haploinsufficiency did not alter mitochondrial content in skeletal muscle (Figure 5A&B and Figure 5 – supplemental figure 1A). High-resolution respirometry experiments revealed that low mitochondrial PE coincides with reduced pyruvate-stimulated *J*O_2_, without affecting OXPHOS protein content per unit of mitochondria (Figure 5C-E and Figure 5 – supplemental figure 1B). Unlike homozygous deletion of PSD, heterozygous knockout of PSD did not promote oxidative stress or mitochondrial electron leak (Figure 5F-H and Figure 5 – supplemental figure 1C). Taken together, these findings are consistent with the notion that low mitochondrial PE is sufficient to drive systemic and skeletal muscle metabolic inflexibility. To delve deeper into the mechanism by which mitochondrial PE abundance facilitates pyruvate metabolism, we performed additional experiments in murine C2C12 myotubes.

**Figure 5.**
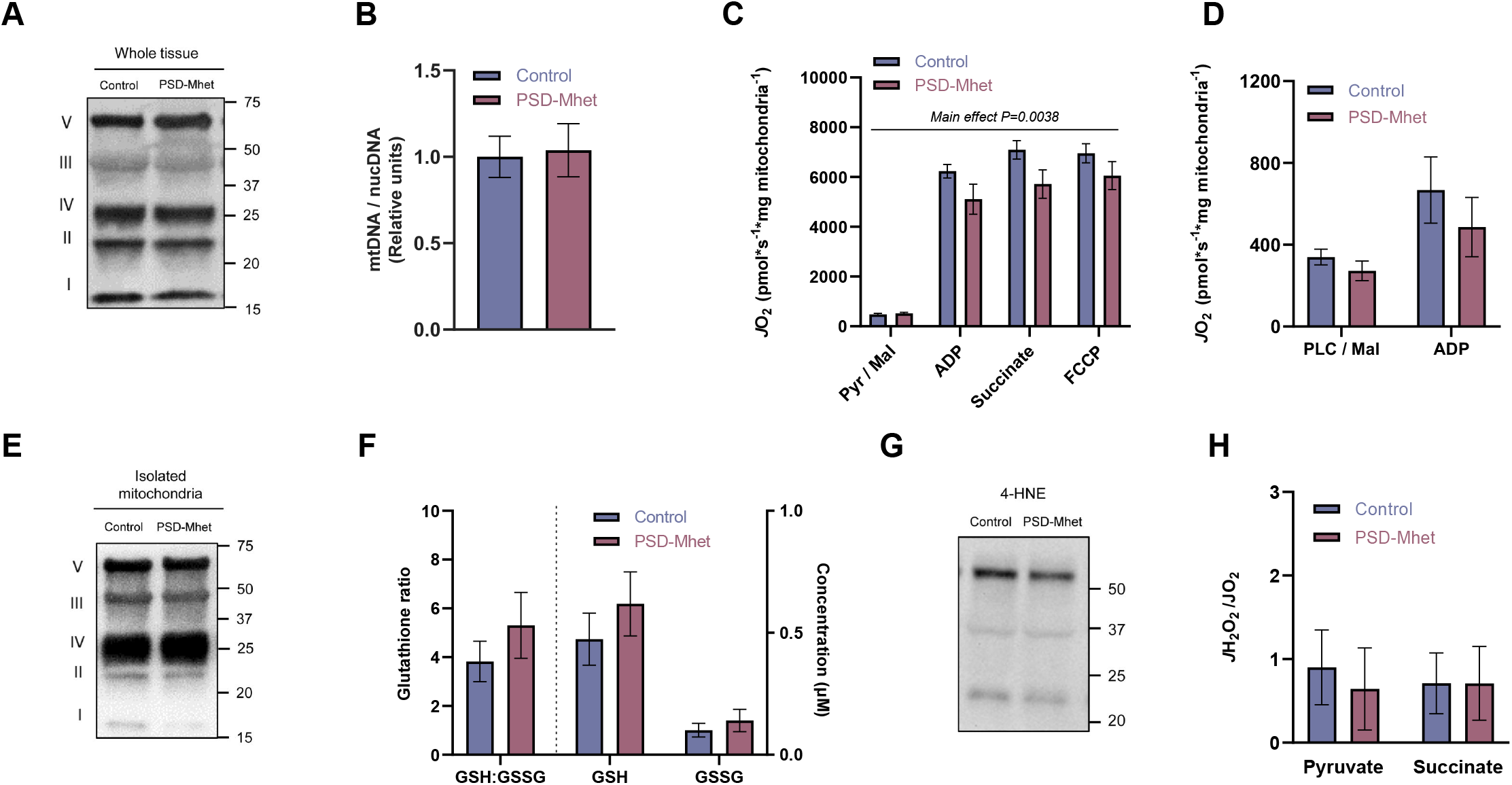
Diminished mitochondrial pyruvate respiration by PSD haploinsufficiency is not mediated by oxidative stress. (A) Representative western blot of ETS protein complexes (I-V) of whole muscle tissue of SMC Control and SMC PSD-MHet mice (n = 4-7 per group). (B) Nuclear to mitochondrial DNA in gastrocnemius muscles (n = 8 per group). (C) O_2_ utilization in isolated muscle mitochondria with TCA cycle substrates using the same conditions described earlier (n = 6 per group). (D) O_2_ utilization in isolated muscle mitochondria with fatty acid substrates using the same conditions described earlier (n = 6-7 per group). (E) Representative western blot of ETS protein complexes (I-V) of isolated muscle mitochondria of SMC Control and SMC PSD-Mhet mice (n = 5 per group). (F) Skeletal muscle glutathione levels (n = 8 per group). (G) Representative 4-HNE western blot of whole muscle of SMC Control and SMC PSD-Mhet mice (n = 6 per group). (H) Electron leak in isolated muscle mitochondria stimulated with succinate or pyruvate and auranofin (n = 6 per group). Data represent mean ± SEM. P-values generated by two-tailed, equal variance, Student’s t-test (B), or by 2-way ANOVA with Tukey’s post hoc test (C-D, F, and H).

### Mitochondrial PE deficiency impairs pyruvate metabolism

To study the effects of low mitochondrial PE, C2C12 myotubes were subjected to lentivirus-mediated knockdown with shRNA encoding either scrambled (shSC) or PSD (shPSD), which was confirmed by qPCR (Figure 6A). We took advantage of the slow turnover rate for phospholipid molecules and performed all experiments 3 days post-lentiviral infection to model modest reductions in mitochondrial PE (Figure 6B). Consistent with our observations *in vivo*, PSD knockdown attenuated pyruvate-stimulated *J*O_2_ or *J*ATP (Figure 6C&D), but not palmitate-stimulated *J*O_2_ (Figure 6E). PSD knockdown also had no effect on OXPHOS content (total cellular or mitochondrial), mitochondrial electron leak, or oxidative stress (Figure 6F and Figure 6 – figure supplement 1A-F). These findings indicate that cell-autonomous effects of PSD deletion are responsible for the phenotype observed *in vivo*.

**Figure 6.**
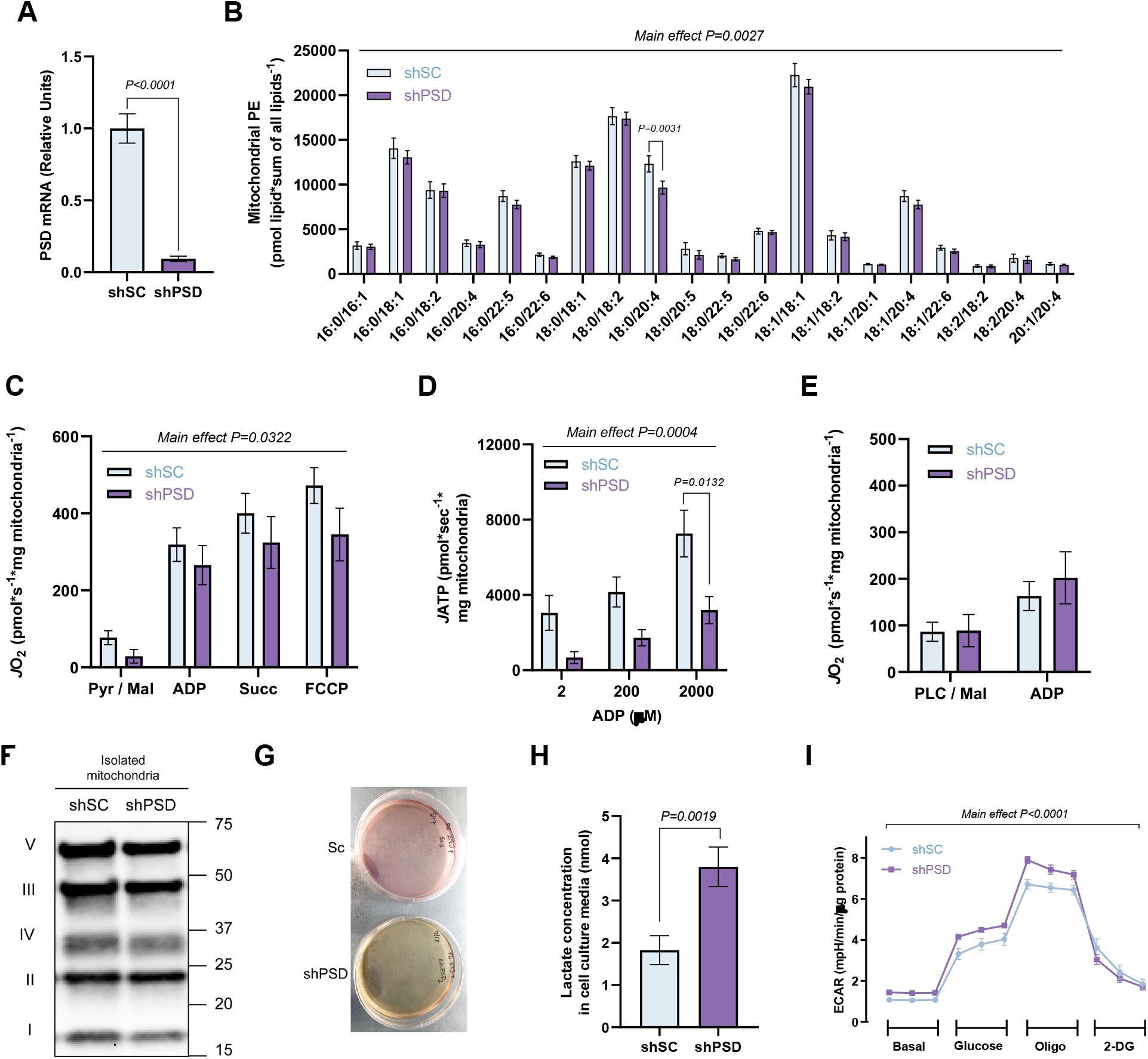
Mitochondrial PE deficiency impairs pyruvate metabolism. (A) PSD mRNA abundance in shSC and shPSD knockdown C2C12 myotubes (n = 9 per group). (B) PE levels from isolated mitochondria from shSC and shPSD cells (n = 9-10 per group). (C) O_2_ consumption with TCA cycle substrates using the same conditions described earlier (n = 7 per group). (D) ATP production in isolated mitochondria from shSc and shPSD myotubes measured in the presence of 0.5 mM malate, 5 mM pyruvate, 5 mM glutamate, 10 mM succinate and either 2, 200, or 2000 μM ADP (n = 7-10 per group). (E) O_2_ consumption with fatty acid substrates using the same conditions described earlier (n = 5-6 per group). (F) Representative western blot of ETS protein complexes I-V in isolated mitochondria from shSC and shPSD cells (n = 5-6 per group). (G) Representative image of media color from cell culture plates. (H) Quantification of lactate production in the media after 24 hours (n = 7-12 per group). (I) Seahorse extracellular acidification rate (ECAR) (n = 14 replicates per group). Data represent mean ± SEM. P-values generated by two-tailed, equal variance, Student’s t-test (A and H), or by 2-way ANOVA with Tukey’s post hoc test (B-E and I).

Knockdown of PSD very strikingly accelerated the yellowing of the culture medium compared to shSC cells (Figure 6G). Yellowing of cell culture media is usually indicative of higher acidification rate due to lactate production [38]. Indeed, lactate concentration in the media was substantially elevated in shPSD cells compared to shSC controls (Figure 6H), and analysis of C2C12 myotubes on the Seahorse Bioanalyzer revealed increased extracellular acidification (ECAR) rate with PSD deletion (Figure 6I). Together, these data likely indicate that low PE causes mitochondria to become resistant to pyruvate metabolism [38, 39].

To more closely examine intracellular pyruvate metabolism, we performed stable isotope tracing using uniformly labeled ^13^C-glucose (Figure 7 and Figure 7 – supplemental figure 1). Targeted mass spectrometry analyses revealed that labeling for glycolytic metabolites leading up to pyruvate was elevated with PSD knockdown (Figure 7B&C), suggesting that low mitochondrial PE does not compromise glucose-to-pyruvate metabolism. Consistent with increased lactate concentration in the media, lactate labeling was higher in shPSD cells compared to shSC (Figure 7D). In contrast, low mitochondrial PE was not associated with increased labeling towards TCA intermediates (Figure 7E-H), suggesting that flux towards lactate, and not TCA cycle, explains the increased labeling for the glycolytic metabolites. These findings are consistent with the notion that mitochondrial PE deficiency impairs mitochondrial pyruvate metabolism.

**Figure 7.**
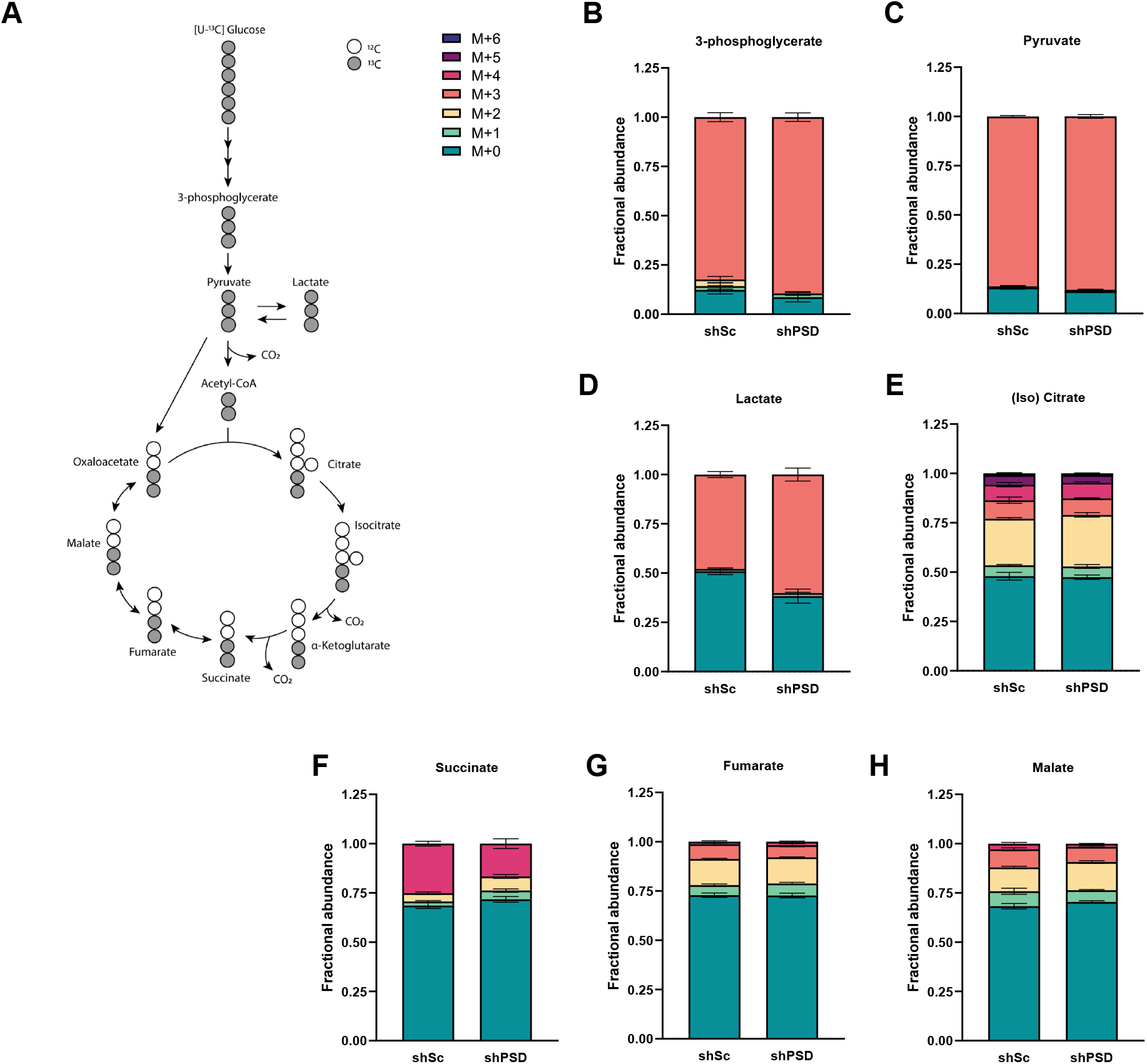
PSD knockdown increases lactate flux. (A) Atom mapping for [U-^13^C_6_]-glucose tracing incorporation into glycolytic and TCA cycle intermediates. White circles represent ^12^C atoms, while black circles signify ^13^C atoms. Isotope labeling pattern between shSC and shPSD myotubes for intracellular (B) 3-phosphoglycerate, (C) pyruvate, (D) lactate, (E) (iso)citrate, (F) succinate, (G) fumarate, and (H) malate (n = 4-5 per group). Data represent mean ± SEM. P-values generated by 2-way ANOVA with Tukey’s post hoc test (B-I).

### Mitochondrial PE facilitates mitochondrial pyruvate entry

We sought to identify the mechanism by which low mitochondrial PE attenuates pyruvate metabolism. Surprisingly, PSD deletion did not reduce protein or mRNA abundance of mitochondrial pyruvate carriers (MPC1 and MPC2) or pyruvate dehydrogenase (PDH) (Figure 8A and Figure 8 – figure supplement 1A&B), suggesting that attenuated pyruvate metabolism is not explained by changes in abundance of these proteins. In fact, there was a statistically significant increase in LDH and a trend for an increase in PDH with PSD deletion. PSD is localized at the inner mitochondrial membrane to generate PE. Thus, we reasoned that the mitochondrial PE may regulate the activity of MPC, which also resides in the inner mitochondrial membrane [40, 41].

**Figure 8.**
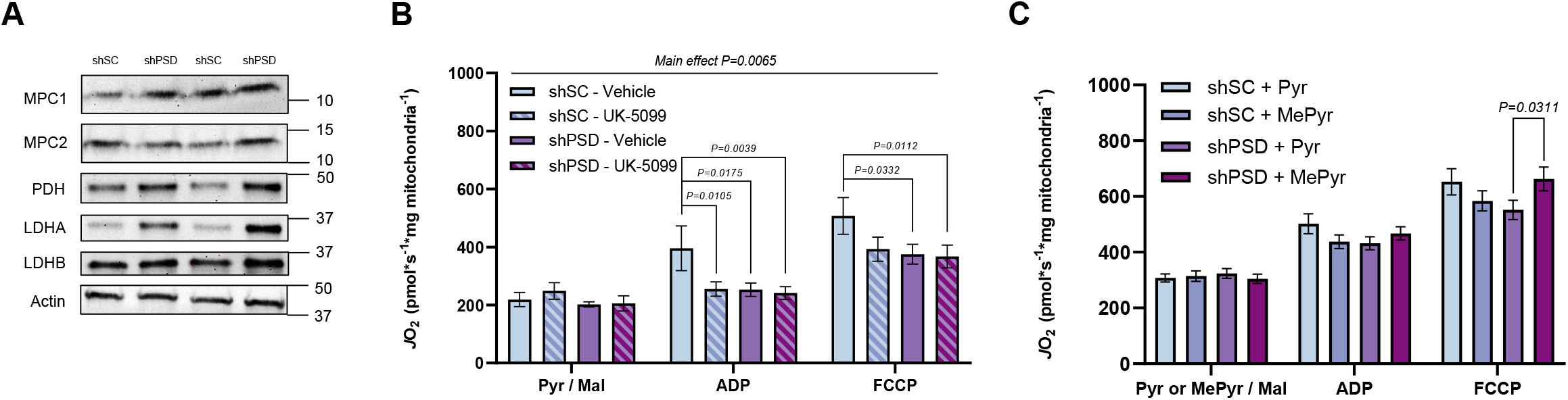
Mitochondrial PE facilitates pyruvate entry. (A) Representative western blot of MPC1, MPC2, PDH, LDHA, LDHB, and Actin between shSC and shPSD myotubes (n = 6 per group). (B) Pyruvate-dependent O_2_ consumption in isolated mitochondria from shSc and shPSDKD myotubes in the presence or absence of the MPC inhibitor, UK-5099 (100nm) and the same Krebs cycle substrate conditions described above (n = 6-8 per group). (C) Pyruvate-dependent respiration in isolated mitochondria with Krebs cycle substrate conditions described above with either pyruvate or methyl pyruvate as a substrate (n = 6-8 per group). Data represent mean ± SEM. P-values generated by 2-way ANOVA with Tukey’s post hoc test (B-C).

To test this possibility, we took a two-pronged approach to link MPC to a defect in pyruvate metabolism. First, we performed pyruvate-stimulated respirometry with or without the MPC inhibitor UK-5099 [42]. Consistent with UK-5099’s action on MPC, pyruvate-stimulated *J*O_2_ was significantly reduced in shSC myotubes (Figure 8B). As expected, MPC inhibition did not completely suppress *J*O_2_ due to anaplerosis. Strikingly, MPC-inhibited *J*O_2_ in shSC cells were similar to *J*O_2_ in shPSD cells without UK-5099, consistent with the notion that reduced *J*O_2_ in shPSD cells is due to attenuated MPC activity. Furthermore, UK-5099 had no effect on *J*O_2_ in shPSD cells, confirming that residual *J*O_2_ in shPSD cells is independent of pyruvate entry via MPC. Second, we compared *J*O_2_ in response to pyruvate or methyl-pyruvate (MePyr). MePyr is a pyruvate-analog that can bypass the MPC, diffuse freely into the mitochondrial matrix, and subsequently demethylated to become mitochondrial pyruvate [43]. MePyr rescued *JO_2_* in shPSD myotubes to pyruvate-stimulated *J*O_2_ levels in shSC cells (Figure 8C). Taken together, these findings suggest that low mitochondrial PE attenuates MPC activity to inhibit mitochondrial pyruvate metabolism.

## Discussion

Skeletal muscle disuse or physical inactivity is linked to 35 chronic diseases [4, 44]. Many of these conditions are attributed to metabolic disturbances caused by sedentary behavior. Nevertheless, the mechanisms by which physical inactivity alters systemic and skeletal muscle metabolism have been poorly defined, likely due to the lack of pre-clinical models [11, 12]. In this study, we developed a novel mouse model of inactivity that reliably induces metabolic inflexibility in male C57BL/6J mice. Metabolic inflexibility was likely driven by pyruvate resistance in skeletal muscle mitochondria. We implicate inactivity-induced downregulation of mitochondrial PE as a driver of pyruvate resistance. Mice with skeletal muscle-specific deletion of PSD was sufficient to recapitulate metabolic inflexibility and mitochondrial pyruvate resistance *in vivo* and *in vitro*. Using stable isotope tracing and high-resolution respirometry, we demonstrate that PE likely directly acts on MPC to facilitate mitochondrial pyruvate entry.

Oxidative stress has been implicated in pathogenesis of inactivity-induced metabolic inflexibility [4, 44]. Indeed, skeletal muscle oxidative stress is commonly manifested in many of the traditional models of mouse disuse [11, 12]. However, while these models are useful in studying muscle atrophy, mice do not develop systemic and skeletal muscle metabolic adaptation observed with human sedentary behavior [11, 12]. In our newly developed SMC model, metabolic inflexibility and suppression of glucose metabolism is faithfully recapitulated, but muscles from this model of inactivity did not exhibit oxidative stress (glutathione, lipid hydroperoxides, mitochondrial electron leak). Notably, our findings from the SMC model reconcile with results from human bedrest studies that oxidative stress cannot explain metabolic inflexibility [36].

Previously we demonstrated that muscle mitochondrial PE becomes elevated with exercise training and decreased with hindlimb unloading [15]. Unlike oxidative stress, SMC reduced skeletal muscle mitochondrial PE concomitant to the development of metabolic inflexibility. What are the mechanisms by which exercise or inactivity promotes changes in muscle mitochondrial PE? In our previous study, as well as in the current study, changes in mitochondrial PE coincided with mRNA abundance of PSD, an enzyme that generates PE in the inner mitochondrial membrane. We believe that changes in PSD levels likely drive the changes in mitochondrial PE abundance. It is currently unknown whether PSD activity is regulated by post-translational modification. It is also possible that there are changes in the upstream mechanism for mitochondrial PE synthesis. PSD generates PE from mitochondrial PS, which is synthesized by PS synthase 1 and 2 in the endoplasmic reticulum [45, 46] and transported to mitochondria via Prelid3b [47]. Finally, it would be important to determine mechanism for the transcriptional control of PSD.

By an unknown reason, PE generated at the endoplasmic reticulum by the Kennedy Pathway do not enter mitochondria [16]. This is exemplified by findings that inhibition of PE synthesis at the ER does not reduce mitochondrial function in skeletal muscle [48, 49]. In fact, deletion of ECT (CTP:phosphoethanolamine cytidylyltransferase, an intermediate step in PE synthesis) increases mitochondrial content, an observation that may be explained by a compensatory increase in muscle PSD [48]. Similarly, deletion of CEPT1 (choline/ethanolamine phosphotransferase, the final step in PE synthesis) increases skeletal muscle glucose metabolism [49]. Thus, combined with our previous report on muscle-specific homozygous deletion of PSD [15], the current study emphasizes that the mitochondrial PE pool remains distinct from that of the endoplasmic reticulum. This is also consistent with findings in yeast, as PE generated by PSD with a forced localization at the outer mitochondrial membrane or endoplasmic reticulum have differential cellular consequences [50].

One of the critical findings of this study was that low mitochondrial PE coincided with pyruvate resistance, but not with palmitate-stimulated *J*O_2_. We demonstrate that PE likely directly facilitates MPC to promote mitochondrial pyruvate uptake, which takes place across the inner mitochondrial membrane where PE is enriched. Meanwhile, the rate-limiting step for fatty acid oxidation is at the step of carnitine palmitoyl transferase-1 (CPT1), which is localized on the outside of the outer mitochondrial membrane [51]. Not only is CPT1 not a transmembrane protein, but it is also localized at the outer mitochondrial membrane where PE is less concentrated [52]. The enzyme equivalent to the MPC for fatty acid oxidation is carnitine/acylcarnitine translocase which is located in the inner mitochondrial membrane, but this enzyme is not the rate-limiting step of palmitate entry nor palmitate oxidation [53, 54]. Thus, we believe that differential subcellular localization of the rate-limiting step for pyruvate or palmitate oxidation contributes to the disproportionate influence of low mitochondrial PE on substrate preference.

Yellowing of cell culture media was the most apparent and robust phenotype observed with PSD knockdown *in vitro*. Our flux experiments reveal that this is a direct result of accelerated flux of glucose towards lactate. Experiments with UK-5099 and MePyr suggest that pyruvate resistance in PSD deficient cells are attributed to the effects of PE on MPC. Multiple studies show that inhibition of MPC promotes resistance for mitochondria to oxidize glycolytic substrates [40, 41, 55, 56]. We believe that the effects of PE deficiency on MPC is the mechanism behind the metabolic inflexible phenotype observed in PSD-MHet mice. We further reason that metabolic inflexibility caused by sedentariness is attributed to low mitochondrial PE which in turn reduces mitochondrial pyruvate entry. It would be important for future studies to elucidate whether PE directly affects the stability of MPC or its post-translational modifications to regulate pyruvate entry.

In conclusion, the current study demonstrates a novel mechanism by which PE facilitates mitochondrial pyruvate entry. We show that a modest reduction in mitochondrial PE is sufficient to promote resistance towards pyruvate oxidation both *in vitro* and *in vivo*. These observations were further extrapolated by findings that pyruvate resistance can be rescued by the membrane permeable MePyr, and that the MPC inhibitor UK-5099 can phenocopy the effects of low mitochondrial PE. We propose that this process drives the metabolic inflexibility induced by physical inactivity. Resistance to pyruvate oxidation may represent a selective advantage for mammals in a state of reduced energy demand, such that substrates are shunted away from skeletal muscle and stored away for subsequent energetic needs. In the modern age of abundant food supply, inactivity-driven resistance for glycolytic substrates can exacerbate the development of metabolic diseases.

## Materials and methods

### Animals

Eight-week old C57BL/6J mice were purchased from the Jackson Laboratory (Strain# 000664) for initial small mouse cage experiments. Heterozygous PSD-MHet mice were generated by crossing our conditional PSD knockout (PSDcKO^+/+^) mice (previously described [15]). PSDcKO^+/+^ mice harbor loxP sites flanking exons 4 to 8 of the mouse PSD gene. These mice were crossed with HSA-MerCreMer mice (HSA-MerCreMer, tamoxifen inducible α-human skeletal actin Cre, courtesy of K. Esser, University of Florida). All mice were bred onto C57BL/6J background and were born at normal Mendelian ratios. Tamoxifen (final concentration of 10 mg ml^-1^) is injected intraperitoneally (7.5μL/g of bodyweight) to PSD-Mhet mice and their respective controls for 5 consecutive days. Mice were maintained on a 12-hour light/12-hour dark cycle in a temperature-controlled room. All animals were fasted for 4 hours prior to tissue collection or experiments. Prior to all terminal experiments and tissue harvesting, mice were given an intraperitoneal injection of 80 mg/kg ketamine and 10 mg/kg xylazine. All protocols were approved by Institutional Animal Care and Use Committee at the University of Utah.

### Small mouse cage

Modified and further developed from Mahmassani et al. [14] and Marmonti et al. [13], SMC is a rectangular box produced from acrylic plastic, made at the University of Utah’s Machine Shop Core. Bedding is placed one-third of the height leaving 4 cm of clearance height. Air holes are designed on all four sides to facilitate air circulation. One air hole on the side was plugged with a Hydropac water lixit (Lab Products Inc., Seaford, Delaware) providing water ad libitum and one air hole on the top is compatible with the hydration system of the Columbus Instruments Oxymax Lab Animal Monitoring System (CLAMS) for determination of whole animal energy expenditure. Abundance of food is provided on top of the bedding to allow ad libitum food consumption. Variable water leakage and crumbling of food are caveats to the attainment of accurate food and water intake in the SMC. Bedding, food, and water were changed every 2-3 days to ensure cleanliness. Two SMC cages can fit in one regular mouse cage. Some experiments were performed with sham or SMC mice housed in pairs, while other experiments were performed with separate cages for sham or SMC mice.

### Indirect Calorimetry

The Columbus Instruments Lab Monitoring System were used to measure VO_2_, RER (respiratory exchange ratio, VCO_2_/VO_2_), food intake, and physical activity (for sham animals only) of sham and SMC mice during Week 7 or 8 of SMC. Mice were individually caged and acclimated for over 24 h in the system before data were collected. Body composition was determined using the Bruker Minispec NMR (Bruker).

### Glucose tolerance test

Intraperitoneal glucose tolerance tests were performed by injection of 1 mg glucose per gram body mass during Week 8 of SMC, at least 3 days prior to sacrifice. Mice were fasted for 4 hours prior glucose injection. Blood glucose was measured before glucose injection and 15, 30, 60, and 120 minutes after injection via tail bleed with a handheld glucometer (Bayer Contour 7151H). In a separate set of experiments, mice were injected with 1 mg glucose per gram body mass, and blood was taken from the facial vein at the 30-minute time point for insulin quantification.

### *Ex vivo* skeletal muscle [^3^H]2-deoxy-D-glucose uptake

*Ex vivo* glucose uptake was measured in soleus muscle as previously described [cite Funai 2013, 2016]. In brief, soleus muscles were dissected and placed in a recovery buffer (KHB with 0.1% BSA, 8 mM glucose, and 2 mM mannitol) at 37°C for 10 minutes. After incubation in recovery buffer, muscles were moved to preincubation buffer (KHB with 0.1% BSA, 2mM sodium pyruvate, and 6 mM mannitol) with or without 200 μU/mL insulin for 15 minutes for soleus and with or without 600 μU/mL insulin for EDL. After preincubation, muscles were placed in incubation buffer (KHB with 0.1% BSA, 9 mM [^14^C]mannitol, 1 mM [^3^H]2-deoxyglucose) with or without 200 μU/mL insulin for 15 minutes. Contralateral muscles were used for basal or insulin-stimulated measurements. After incubation, muscles were blotted dry on ice-cold filter paper, snap-frozen, and stored at −80°C until analyzed with liquid scintillation counting.

### Serum insulin, glucose, and cortisol quantification

Blood was collected from facial vein either before anesthesia or at the 30-minute time point of the glucose tolerance test. Blood was then placed at room temperature for 20 minutes to clot prior to centrifugation at 2000 x g for 10 minutes at 4°C. The supernatant (serum) was collected, placed in a new tube, and stored at until −80°C analysis. Serum insulin levels were quantified using a Mouse Insulin ELISA kit (Cat# 90080 Crystal Chem, Chicago, Illinois). Serum glucose was quantified using a Mouse Glucose Assay Kit (Cat# 81692 Crystal Chem, Chicago, Illinois). Serum cortisol levels were quantified by the DetectX ELISA kit (Cat# K003-H1W Arbor assays, Chicago, USA).

### High-resolution respirometry and fluorometry

Mitochondrial O_2_ utilization was measured using the Oroboros O2K Oxygraphs. Skeletal muscle tissues were minced in mitochondria isolation medium (300 mM sucrose, 10 mM HEPES, 1 mM EGTA) and subsequently homogenized using a Teflon-glass system. Homogenates were then centrifuged at 800 x g for 10 min, after which the supernatant was taken and centrifuged at 12,000 x g for 10 min. The resulting pellet was carefully resuspended in mitochondria isolation medium. Isolated mitochondria were then added to the oxygraphy chambers containing assay buffer (MES potassium salt 105 mM, KCl 30 mM, KH_2_PO_4_ 10 mM, MgCl2 5 mM, BSA 0.5 mg/ml). Respiration was measured in response to the following substrates: 0.5mM malate, 5mM pyruvate, 5mM glutamate, 10mM succinate, 1.5 μM FCCP, 0.02mM palmitoyl-carnitine, 5mML-carnitine. ATP production was measured fluorometrically using a Horiba Fluoromax 4 (Horiba Scientific), by enzymatically coupling ATP production to NADPH synthesis as previously described [57]. Respiration and ATP production were measured in the presence of 2mM ADP, unless otherwise stated.

For inhibitor experiments in mitochondria isolated from shSC and shPSD myotubes, the mitochondrial pyruvate carrier (MPC) inhibitor, UK-5099 (5048170001, Sigma Aldrich), was used to inhibit MPC activity. To induce a submaximal drop of pyruvate-dependent respiration, 100 nM UK-5099 was used at a submaximal concentration and injected into the oxygraph chamber following the addition of malate and pyruvate. Respiration was measured in response to the following substrates: 0.5 mM malate, 5 mM pyruvate, 2 mM ADP, and 1 μM FCCP. To evaluate whether pyruvate-dependent respiration was compromised in shSC and shPSD mitochondria, respiration was measured in response to either 5 mM pyruvate or 5 mM methyl pyruvate (371173, Sigma Aldrich) along with the above substrates.

### H_2_O_2_ measurements

Mitochondrial H_2_O_2_ emission was determined in isolated mitochondria from skeletal muscle and permeabilized muscle fibers. All *J*H_2_O_2_ experiments were performed in buffer Z supplemented with 10 mM Amplex UltraRed (Invitrogen), 20 U/mL CuZn SOD, and 25 mM Blebbistatin (for permeabilized muscle fibers only). Briefly, isolated mitochondria or permeabilized fibers were added to 1 ml of assay buffer containing Amplex Ultra Red, which produces a fluorescent product when oxidized by H_2_O_2_. H_2_O_2_ emission was measured following the addition of 10mM succinate or 5 mM pyruvate for a final concentration. The appearance of the fluorescent product was measured by a Horiba Fluoromax 4 fluorometer (excitation/emission 565/600).

### Seahorse assay

Extracellular acidification rate (ECAR) was measured in C2C12 myoblasts using a Seahorse XF96 Analyzer. Myoblasts were plated at 5 x 10 cells/well and grown in lentiviral media for 48 hours. C2C12 cells were selected with puromycin throughout differentiation for 3 days. The real-time extracellular acidification rate (ECAR) was measured using the XFe96 extracellular flux analyzer with the Glycolysis Stress Kit (Agilent Technologies) following the manufacturer’s instructions. The measurement was normalized to total protein determined by Pierce BCA Protein Assay Kit (ThermoFisher). Briefly, cells were seeded on XF96 cell culture microplates (Seahorse Bioscience) at a seeding density of 5.0 x 10^3^ cells per well. Before assay, cells were rinsed twice and kept in pre-warmed XF basic assay medium (pH 7.4) supplemented with 2 mM glutamine in a 37°C non-CO_2_ incubator for an hour. Then the rate was measured at 37°C in 14 replicates (separate wells) by using the following compounds in succession: 10 mM glucose, 1 μM oligomycin, and 50 mM 2-DG. Basal ECAR was measured before drug exposure. The glycolytic function metrics was calculated by Seahorse Wave Desktop Software as directed in the glycolysis stress kit manual (Agilent Technologies).

### Glutathione Redox

Skeletal muscle GSH and GSSG was measured using the fluorometric GSH/GSSG Ratio Detection Assay Kit II (Abcam 205811). Briefly, whole muscle was homogenized in lysis buffer, deproteinized using the Deproteinizing Sample Kit – TCA (Abcam 204708), nutated at 4°C for 1 hour, and centrifuged at 4°C for 15 min at 12,000*g*. The supernatant was collected and protein concentrations were determined using the Pierce BCA Protein Assay (Thermo Fischer Scientific). Supernatant was then used to determine GSH and total glutathione. Fluorescence was measured at Ex/Em = 490/520nm with a fluorescence microplate reader.

### Cell culture

C2C12 myoblasts were grown in high-glucose DMEM (4.5 g/L glucose, with L-glutamine; Gibco 11965-092) supplemented with 10% FBS (heat-inactivated, certified, US origin; Gibco 10082-147), and 0.1% penicillin-streptomycin (10,000 U/mL; Gibco 15140122). C2C12 cells were differentiated into myotubes with low-glucose DMEM (1 g/L glucose with 4mM L-glutamine and 110 mg/L sodium pyruvate; Gibco 11885-084) supplemented with 2% horse serum (defined; VWR 16777), and 0.1% penicillin-streptomycin.

### Lentivirus-mediated knockdown of PSD

PSD expression was decreased using pLKO.1 lentiviral-RNAi system. Plasmids encoding shRNA for mouse PISD (shPSD: TRCN0000115415) was obtained from MilliporeSigma. Packaging vector psPAX2 (ID 12260), envelope vector pMD2.G (ID 12259), and scrambled shRNA plasmid (SC: ID 1864) were obtained from Addgene. HEK293T cells in 10 cm dishes were transfected using 50 μL 0.1% polyethylenimine, 200 μL, 0.15 M sodium chloride, and 500 μL Opti-MEM (with HEPES, 2.4 g/L sodium bicarbonate, and l-glutamine; Gibco 31985) with 2.66 μg of psPAX2, 0.75 μg of pMD2.G, and 3 μg of either scrambled or PISD shRNA plasmid. Cells were selected with puromycin throughout differentiation to ensure that only cells infected with shRNA vectors were viable.

### U-^13^C glucose labeling in cultured myotubes

For metabolite extraction, cold 90% methanol (MeOH) solution was added to each sample to give a final concentration of 80% MeOH to each cell pellet. Samples were then incubated at −20 °C for 1 hr. After incubation, samples were centrifuged at 20,000 x g for 10 minutes at 4 °C. The supernatant was then transferred from each sample tube into a labeled, fresh micro centrifuge tube. Process blanks were made using only extraction solvent and no cell culture. The samples were then dried *en vacuo*.

All GC-MS analysis was performed with an Agilent 5977b HES fit with an Agilent 7693A automatic liquid sampler. Dried samples were suspended in 40 μL of a 40 mg/mL O-methoxylamine hydrochloride (MOX) (MP Bio #155405) in dry pyridine (EMD Millipore #PX2012-7) and incubated for one hour at 37 °C in a sand bath. 25 μL of this solution was added to auto sampler vials. 60 μL of N-methyl-N-trimethylsilyltrifluoracetamide (MSTFA with 1%TMCS, ThermoFisher Scientific #TS48913) was added automatically via the auto sampler and incubated for 30 minutes at 37 °C. After incubation, samples were vortexed and 1 μL of the prepared sample was injected into the gas chromatograph inlet in the split mode with the inlet temperature held at 250 °C. A 10:1 split ratio was used for analysis for the majority of metabolites. Any metabolites that saturated the instrument at the 10:1 split were analyzed at a 50:1 split ratio. The gas chromatograph had an initial temperature of 60 °C for one minute followed by a 10 °C/min ramp to 325 °C and a hold time of 10 minutes. A 30-meter Agilent Zorbax DB-5MS with 10 m Duraguard capillary column was employed for chromatographic separation. Helium was used as the carrier gas at a rate of 1 mL/min.

Data was collected using the Agilent MassHunter software. Metabolites were identified and their peak area was recorded using MassHunter Quant. Metabolite identity was established using a combination of an in-house metabolite library developed using pure purchased standards and the NIST and Fiehn libraries. There are a few reasons a specific metabolite may not be observable through GC-MS. The metabolite may not be amenable to GC-MS due to its size, or a quaternary amine such as carnitine, or simply because it does not ionize well.

### Lipid Extraction

LC-MS-grade solvents and mobile phase modifiers were obtained from Honeywell Burdick & Jackson, Morristown, NJ (acetonitrile, isopropanol, formic acid), Fisher Scientific, Waltham, MA (methyl *tert*-butyl ether) and Sigma–Aldrich/Fluka, St. Louis, MO (ammonium formate, ammonium acetate). Lipid standards were obtained from Avanti Polar Lipids, Alabaster, AL. Lipids were extracted from mitochondria using a modified Matyash lipid extraction [58] using a biphasic solvent system of cold methanol, methyl *tert*-butyl ether (MTBE), and water. Briefly, a mixture of cold MTBE, methanol, and internal standards (Mouse SPLASH LIPIDOMIX Avanti Polar Lipids 330707 and Cardiolipin Mix I Avanti Polar Lipids LM6003) were added to isolated skeletal muscle mitochondria isolated mitochondria from C2C12 myotubes or plantaris skeletal muscle. Samples were sonicated for 60 sec, then incubated on ice with occasional vortexing for 1 hr. After addition of 188 μL of PBS, the mixture was incubated on ice for 15 min and centrifuged at 12,000 x *g* for 10 minutes at 4 °C. The organic (upper) layer was collected, and the aqueous layer was re-extracted with 1 mL of 10:3:2.5 (*v/v/v*) MTBE/MeOH/water. The MTBE layers were combined for untargeted lipidomic analysis and dried under vacuum. The aqueous layer was centrifuged for 12,000 x *g* for 10 minutes at 4 °C and dried under vacuum. Lipid extracts were reconstituted in 500 μL of 8:2:2 (*v/v/v*) IPA/ACN/water containing 10 mM ammonium formate and 0.1% formic acid. Concurrently, a process blank sample was prepared and then a pooled quality control (QC) sample was prepared by taking equal volumes (~50 μL) from each sample after final resuspension.

### LC-MS Analysis and Data Processing

Lipid extracts were separated on an Acquity UPLC CSH C18 column (2.1 x 100 mm; 1.7 μm) coupled to an Acquity UPLC CSH C18 VanGuard precolumn (5 × 2.1 mm; 1.7 μm) (Waters, Milford, MA) maintained at 65 °C connected to an Agilent HiP 1290 Sampler, Agilent 1290 Infinity pump, and Agilent 6545 Accurate Mass Q-TOF dual AJS-ESI mass spectrometer (Agilent Technologies, Santa Clara, CA). Samples were analyzed in a randomized order in both positive and negative ionization modes in separate experiments acquiring with the scan range m/z 100 – 1700. Mobile phase A consisted of ACN:H_2_O (60:40, *v/v*) in 10 mM ammonium formate and 0.1% formic acid, and mobile phase B consisted of IPA:ACN:H_2_O (90:9:1, *v/v/v*) in 10 mM ammonium formate and 0.1% formic acid. For negative mode analysis the modifiers were changed to 10 mM ammonium acetate. The chromatography gradient for both positive and negative modes started at 15% mobile phase B then increased to 30% B over 2.4 min, it then increased to 48% B from 2.4 – 3.0 min, then increased to 82% B from 3 – 13.2 min, then increased to 99% B from 13.2 – 13.8 min where it’s held until 16.7 min and then returned to the initial conditions and equilibriated for 5 min. Flow was 0.4 mL/min throughout, with injection volumes of 2 μL for positive and 10 μL negative mode. Tandem mass spectrometry was conducted using iterative exclusion, the same LC gradient at collision energies of 20 V and 27.5 V in positive and negative modes, respectively. For data processing, Agilent MassHunter (MH) Workstation and software packages MH Qualitiative and MH Quantitative were used. The pooled QC (n=8) and process blank (n=4) were injected throughout the sample queue to ensure the reliability of acquired lipidomics data. For lipid annotation, accurate mass and MS/MS matching was used with the Agilent Lipid Annotator library and LipidMatch [59]. Results from the positive and negative ionization modes from Lipid Annotator were merged based on the class of lipid identified. Data exported from MH Quantitative was evaluated using Excel where initial lipid targets are parsed based on the following criteria. Only lipids with relative standard deviations (RSD) less than 30% in QC samples are used for data analysis. Additionally, only lipids with background AUC counts in process blanks that are less than 30% of QC are used for data analysis. The parsed excel data tables are normalized based on the ratio to class-specific internal standards, then to tissue mass and sum prior to statistical analysis.

### Western blotting

Tissues or cells were homogenized in lysis buffer, nutated at 4°C for 1 hour, and centrifuged at 4°C for 15 min at 12,000*g*, and the supernatant was transferred to a new tube. Western blotting was performed as previously described [60], and samples were analyzed for protein abundance of OXPHOS (ab110413, Abcam), 4-HNE (ab46545, Abcam), MPC1 (generated by Jared Rutter), MPC2 (generated by Jared Rutter), PDH (3205S, Cell Signaling), LDHA (3582S, Cell Signaling), LDHB (sc-100775, SantaCruz Biotech), and Actin (A2066, Sigma-Aldrich).

### RNA quantification

For quantitative polymerase chain reaction (qPCR) experiments, mouse tissues or cells were lysed in the TRIzol reagent (Thermo Fisher Scientific), and RNA was isolated using standard techniques. The iScript cDNA Synthesis Kit was used to reverse transcribe total RNA, and qPCR was performed with SYBR Green reagents (Thermo Fisher Scientific). Pre-validated primer sequences were obtained from mouse primer depot (https://mouseprimerdepot.nci.nih.gov/). All mRNA levels were normalized to RPL32. For RNA sequencing, gastrocnemius muscle RNA from Sham and SMC mice were isolated with the Direct-zol RNA Miniprep Plus kit (Zymo Cat#: R2070). RNA library construction and sequencing were performed by the High-Throughput Genomics Core at the Huntsman Cancer Institute, University of Utah. RNA libraries were constructed using the NEBNext Ultra II Directional RNA Library Prep with rRNA Depletion Kit (human, mouse rat) and the following adapter reads: Read 1: AGATCGGAAGAGCACACGTCTGAACTCCAGTCA and Read 2: AGATCGGAAGAGCGTCGTGTAGGGAAAGAGTGT. Sequencing was performed using the NovaSeq S4 Reagent Kit v1.5 150×150 bp Sequencing with >25 million reads per sample. Pathway analyses were performed by the Bioinformatics Core at the Huntsman Cancer Institute, University of Utah using the KEGG (Kyoto Encyclopedia of Genes and Genomes) Pathway Database. For differentially expressed genes, only transcripts with *P*adj < 0.05 and baseMean > 100 are included.

### DNA isolation and quantitative PCR

Genomic DNA for assessments of mitochondrial DNA (mtDNA) was isolated using a commercially available kit according to the manufacturer’s instructions (69504, Qiagen). Genomic DNA was added to a mixture of SYBR Green (Thermo Fisher Scientific) and primers. Sample mixtures were pipetted onto a 3840well plate and analyzed with QuantStudio 12K Flex (Life Technologies). The following primers were used: mtDNA fwd, TTAAGA-CAC-CTT-GCC-TAG-CCACAC; mtDNA rev, CGG-TGG-CTG-GCA-CGA-AAT-T; nucDNA fwd, ATGACG-ATA-TCG-CTG-CGC-TG; nucDNA rev, TCA-CTT-ACC-TGGTGCCTA-GGG-C.

### Statistical analyses

All data presented herein are expressed as mean ± SEM. The level of significance was set at p < 0.05. Student’s t-tests were used to determine the significance between experimental groups and two-way ANOVA analysis followed by Tukey’s HSD post hoc test was used where appropriate. The sample size (n) for each experiment is shown in the figure legends and corresponds to the sample derived from the individual mice or for cell culture experiments on an individual batch of cells. Unless otherwise stated, statistical analyses were performed using GraphPad Prism software.

## Acknowledgements

This research is supported by National Institutes of Health grants DK107397, DK127979, GM144613, AG074535, AG067186 (to K.F.), DK091317 (to M.J.L.), DK104998 (to E.B.T.), AG076075, AG079477, AG050781 (to M.J.D.), GM131854, CA228346 (to J.R.), Howard Hughes Medical Institute (J.R.), and American Heart Association grant 915674 (to P.S.). University of Utah Metabolomics Core Facility is supported by S10 OD016232, S10 OD021505, and U54 DK110858.

## Author contributions

Piyarat Siripoksup, Conceptualization, Data curation, Formal analysis, Validation, Investigation, Visualization, Methodology, Writing – original draft, Writing – review and editing, Funding acquisition; Guoshen Cao, Conceptualization, Investigation, Data curation, Methodology; Ahmad A Cluntun, Conceptualization, Data curation, Formal analysis, Methodology; J Alan Maschek, Data curation, Formal analysis, Resources; Quentinn Pearce, Data curation, Formal analysis, Resources; Marisa J Lang, Data curation; Hiroaki Eshima, Data curation; Precious C Opurum, Data curation; Ziad S Mahmassani, Conceptualization, Methodology; Eric B Taylor, Methodology; James E Cox, Methodology, Resources, Supervision; Micah J Drummond, Conceptualization, Methodology, Resources, Supervision; Jared Rutter, Conceptualization, Resources; Katsuhiko Funai, Conceptualization, Formal analysis, Validation, Visualization, Supervision, Writing – review and editing, Resources, Funding acquisition, Project administration.

## Ethics

Experiments on animals were performed in strict accordance with the Guide for the Care and Use of Laboratory Animals of the National Institutes of Health. All animals were handled according to approved University of Utah Animal Use and Care Committee (IACUC) protocols (#20-07007). The protocol as approved by the Committee on the Ethics of Animal Experiments of the University of Utah.

---

**Figure 1 – Figure Supplement 1.**
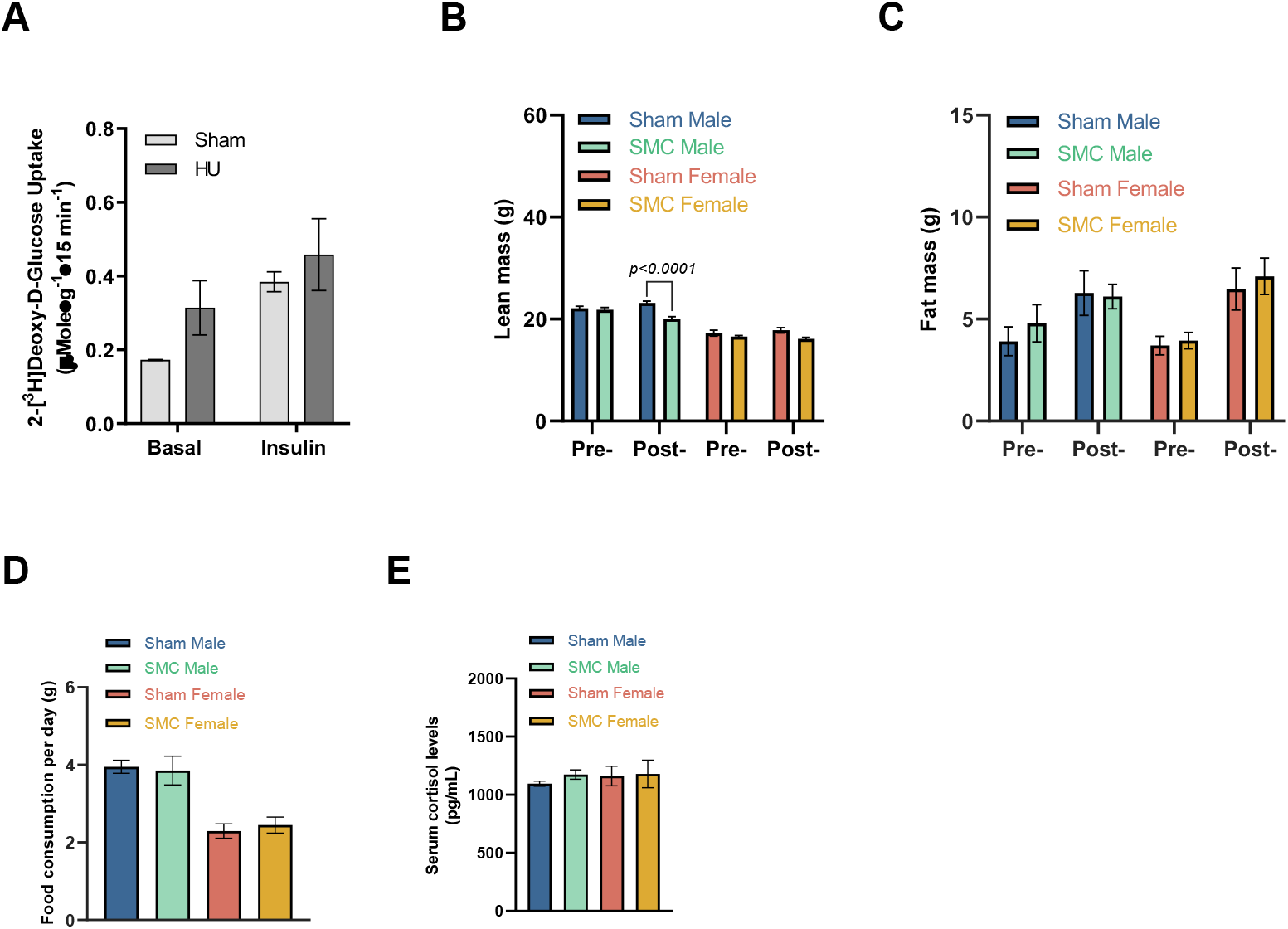
SMC housing induces metabolic inflexibility in male but not female mice. (A) [^3^H]2-deoxyglucose glucose uptake in soleus muscles of sham and hindlimb unloading (HU) mice (n = 2 per group). (B) Lean mass pre- and post-intervention by NMR (n = 12-15 per group). (C) Fat mass pre- and post-intervention by NMR (n = 4-8 per group). (D) Food consumption (n = 18-28 per group). (E) Fasting serum cortisol levels (n = 6-7 per group). Data represent mean ± SEM. P-values generated by two-tailed, equal variance, Student’s t-test (B, D, E), or by 2-way ANOVA with Tukey’s post hoc test (A, C).

**Figure 2 – Figure Supplement 1.**
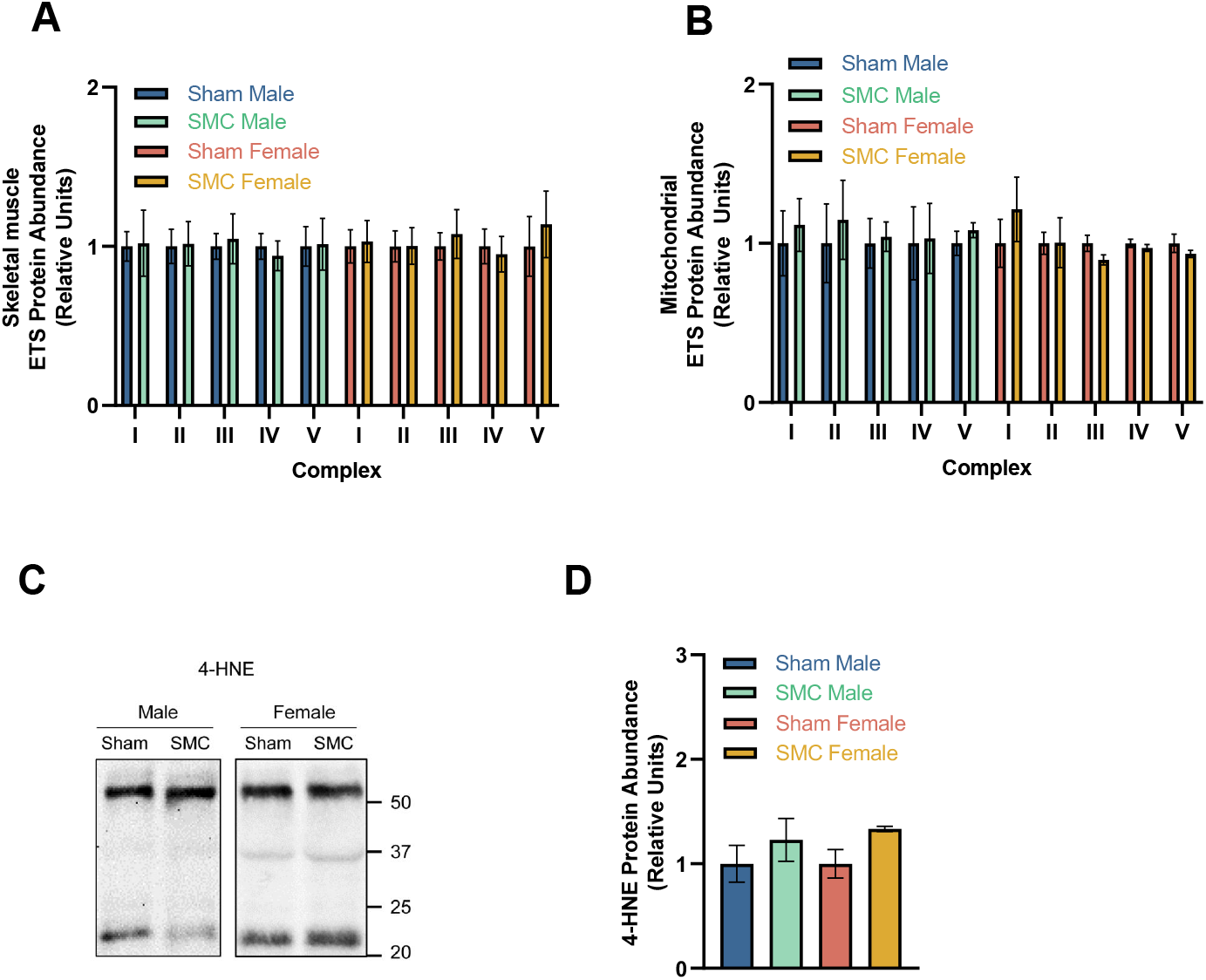
SMC housing reduces pyruvate-dependent respiration without altering palmitate-stimulated respiration. (A) Skeletal muscle ETS protein quantification (n = 3-4 per group). (B) Isolated muscle mitochondria ETS protein quantification (n = 5-6 per group). (C) Representative 4-hydroxynonenal (4-HNE) western blot of whole muscle tissue of sham and SMC mice (n = 5-7 per group). (D) 4-HNE protein quantification (n = 5-7 per group). Data represent mean ± SEM. P-values generated by 2-way ANOVA with Tukey’s post hoc test (A-B and D).

**Figure 3 – Figure Supplement 1.**
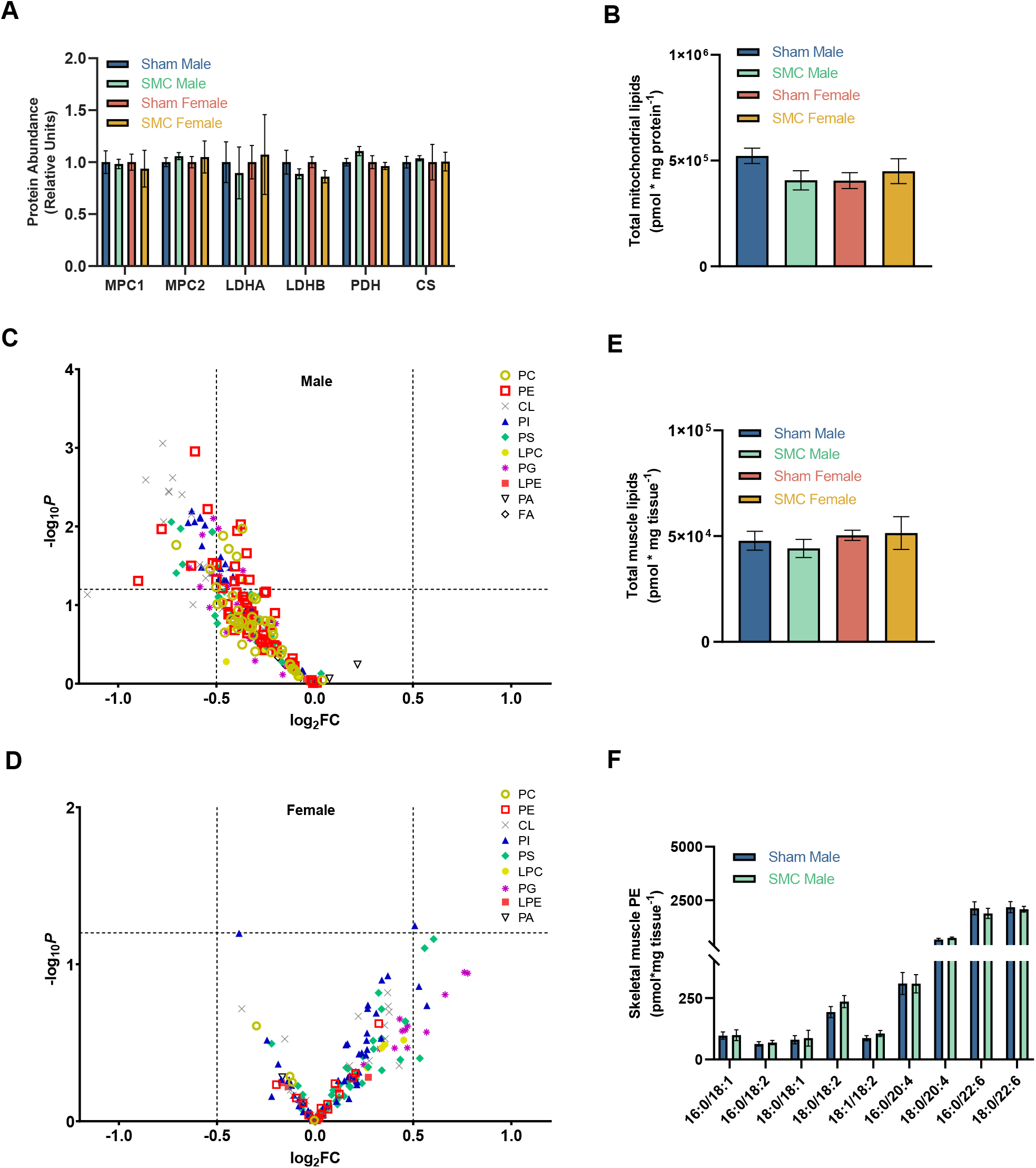
Physical inactivity by SMC housing alters skeletal muscle membrane lipid composition. (A) Western blot quantification of glycolytic/TCA protein abundances in sham and SMC mice (n = 2-6 per group). (B) Total mitochondrial and (E) skeletal muscle lipids between male sham and SMC mice (n = 7-8 per group) and female sham and SMC mice (n = 5 per group). Volcano plot showing changes in muscle mitochondrial lipids between male (C) and female (D) sham and SMC mice (n = 8 per group). (F) Skeletal muscle PE abundance of male sham and SMC mice (n = 7-8 per group). Data represent mean ± SEM. P-values generated by 2-way ANOVA with Tukey’s post hoc test (A-B and E-F).

**Figure 4 – Figure Supplement 1.**
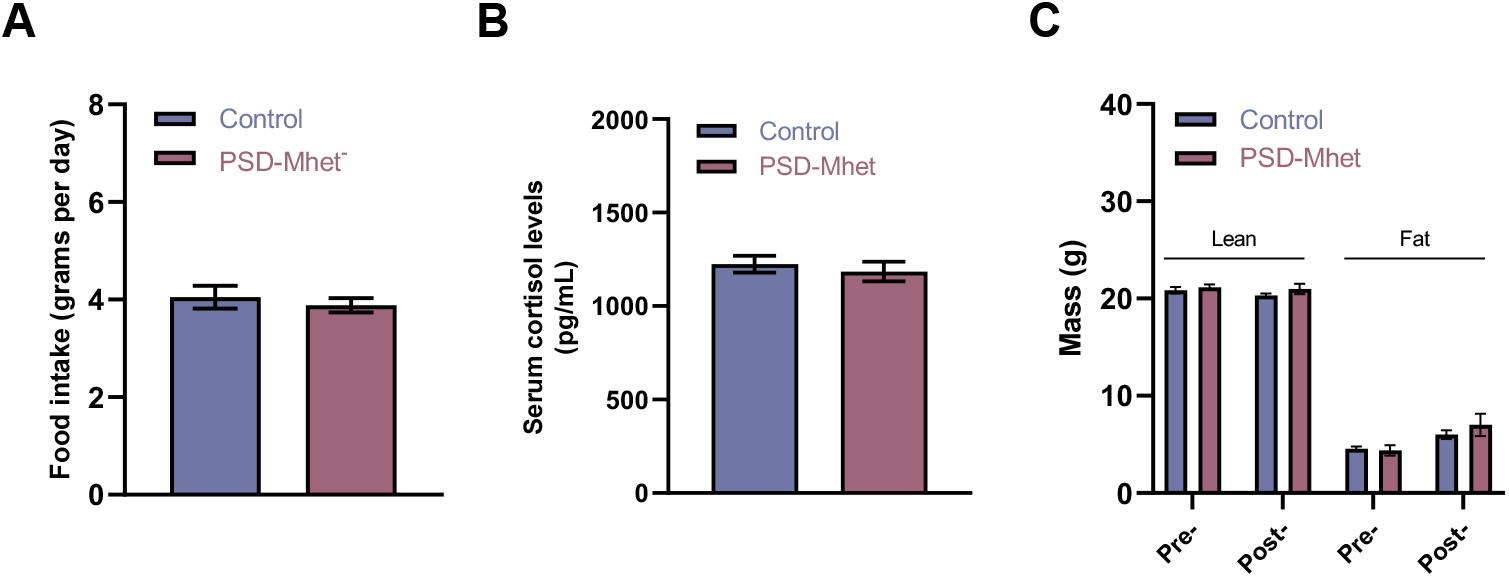
Muscle PSD haploinsufficiency increases susceptibility of mice to inactivity-induced metabolic inflexibility. (A) Average food intake per day throughout SMC intervention (n = 10 per group). (B) Serum cortisol levels after SMC intervention (n = 8 per group). (C) Lean and fat mass by NMR of SMC Control and SMC PSD-Mhet mice pre- and post-intervention (n = 8-12 per group). Data represent mean ± SEM. P-values generated by two-tailed, equal variance, Student’s t-test (A-B), or by 2-way ANOVA with Tukey’s post hoc test (C).

**Figure 5 – Figure Supplement 1.**
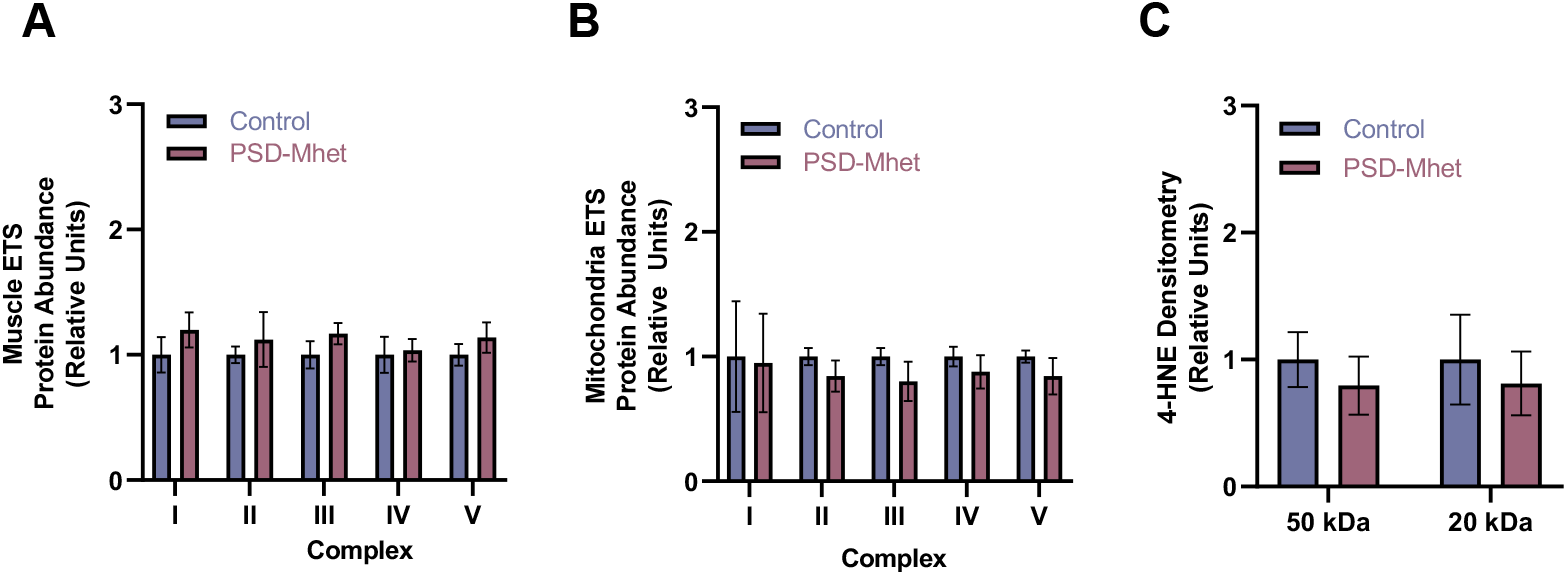
Diminished mitochondrial pyruvate respiration by PSD haploinsufficiency is not mediated by oxidative stress. (A) Whole tissue ETS protein quantification (n = 4-7 per group). (B) Mitochondrial ETS protein quantification (n = 5 per group). (C) 4-HNE protein quantification (n = 6 per group). Data represent mean ± SEM. P-values generated by 2-way ANOVA with Tukey’s post hoc test (A-C).

**Figure 6 – Figure Supplement 1.**
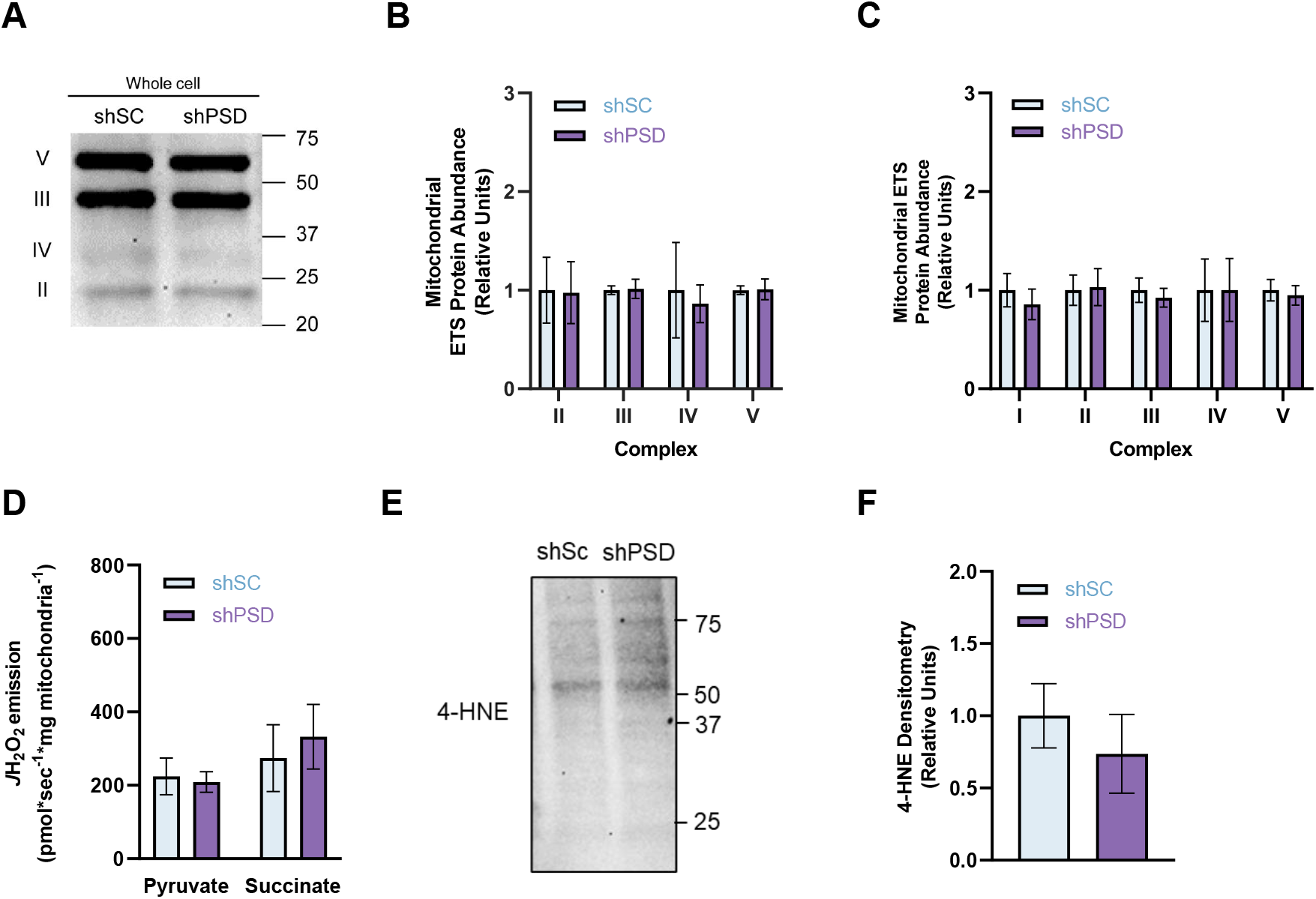
Mitochondrial PE deficiency impairs pyruvate metabolism. (A) Representative western blot of whole cell OXPHOS complexes between shSC and shPSD cells (n = 3-4 per group). (B) Mitochondrial ETS protein quantification (n = 5-6 per group). (C) Quantification of (A) (n = 3-4 per group). (D) H_2_O_2_ emission in isolated mitochondria from shSC and shPSD myotubes stimulated with succinate or pyruvate and auranofin (n = 4-10 per group). (E) Representative western blot of whole cell 4-HNE (n = 6 per group). (F) Quantification of (E) (n = 6 per group). Data represent mean ± SEM. P-values generated by two-tailed, equal variance, Student’s t-test (F) or by 2-way ANOVA with Tukey’s post hoc test (B-D).

**Figure 7 – Figure Supplement 1.**
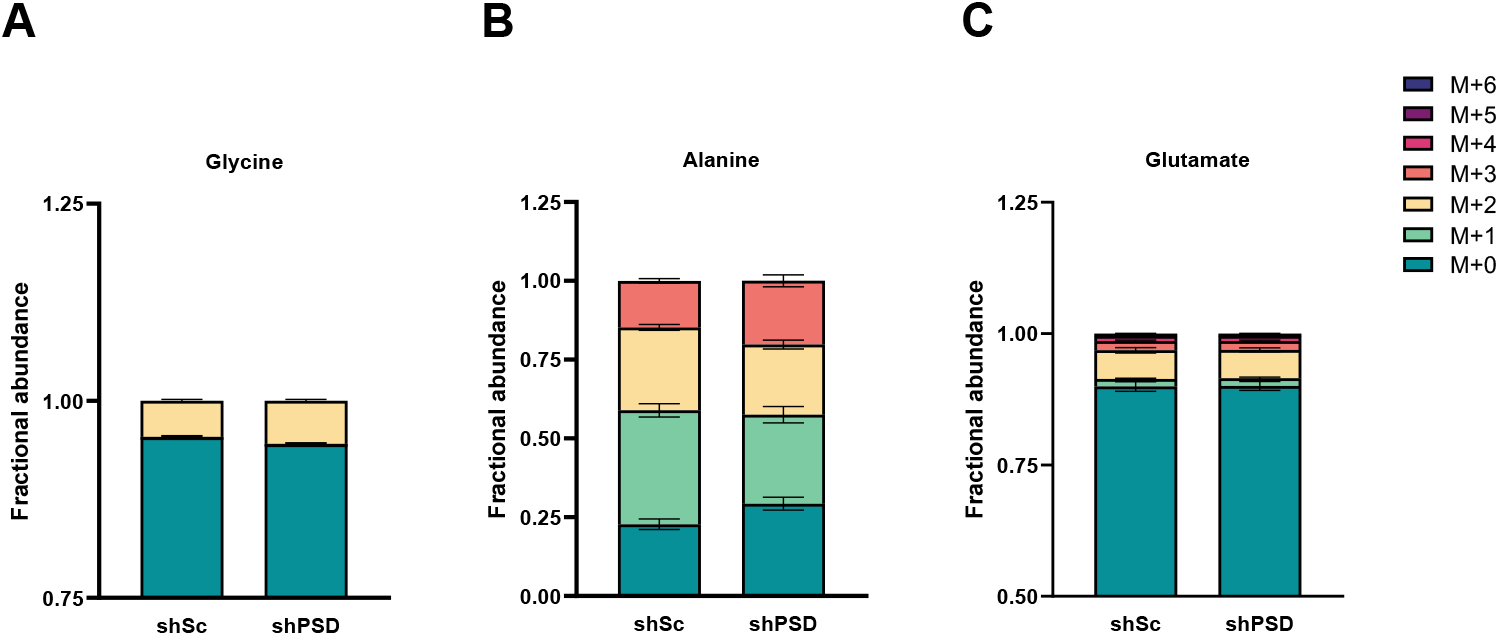
PSD knockdown increases lactate flux. Isotopic labeling pattern in shSC and shPSD myotubes of (A) glycine, (B) alanine, (C) glutamate (n = 4-5 per group). Data represent mean ± SEM. P-values generated by 2-way ANOVA with Tukey’s post hoc test (A-C).

**Figure 8 – Figure Supplement 1.**
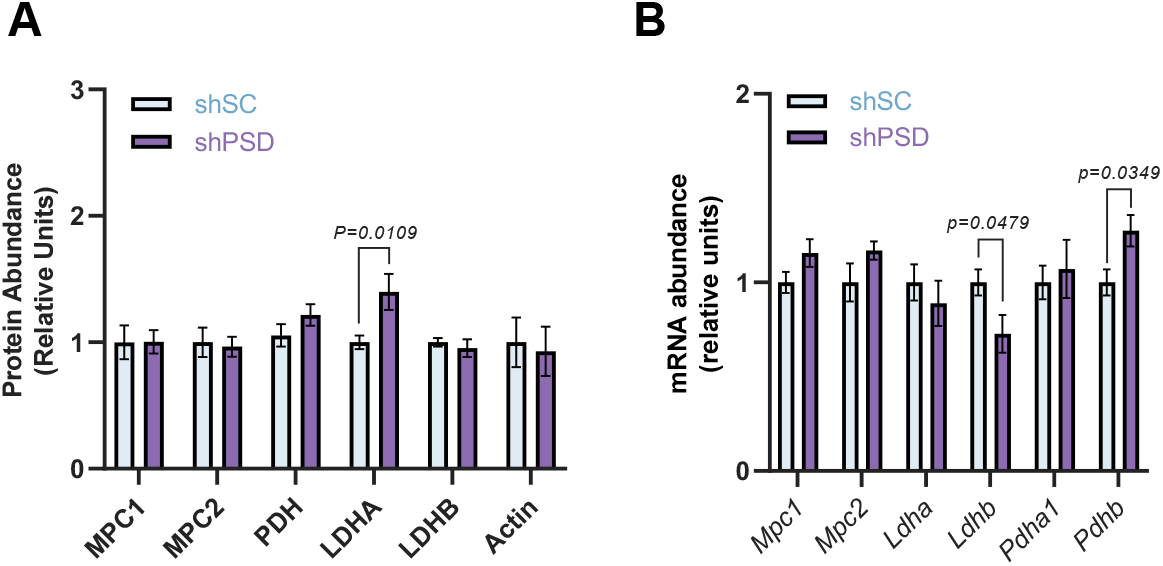
Mitochondrial PE facilitates pyruvate entry. (A) Western blot quantification of MPC1, MPC2, PDH, LDHA, LDHB, and Actin protein abundances (n = 4-6 per group). (B) Relative mRNA abundances of genes encoding for proteins in (A) (n = 6 per group). Data represent mean ± SEM. by two-tailed, equal variance, Student’s t-test (A-B)

